# Disrupted transcriptional networks regulated by *CHD1L* during neurodevelopment underlie the mirrored neuroanatomical and growth phenotypes of the 1q21.1 copy number variant

**DOI:** 10.1101/2025.02.18.638841

**Authors:** Marianne Victoria Lemée, Maria Nicla Loviglio, Tao Ye, Peggy Tilly, Céline Keime, Chantal Weber, Anastasiya Petrova, Pernelle Klein, Bastien Morlet, Olivia Wendling, Hugues Jacobs, Mylène Tharreau, David Geneviève, Juliette D Godin, Christophe Romier, Delphine Duteil, Christelle Golzio

## Abstract

Distal 1q21.1 deletions and duplications are associated with variable phenotypes including autism, head circumference and height defects. To elucidate which gene(s) are responsible for the 1q21.1 duplication/deletion-associated phenotypes, we performed gene manipulation in zebrafish and mice. We modeled 1q21.1 duplication by overexpressing the eight human protein-coding genes in zebrafish. We found that overexpression of *CHD1L* only led to macrocephaly and increased larval body length, whereas chd1l deletion caused opposite phenotypes. These mirrored phenotypes were also observed in mouse embryos. Transcriptomic, cistromic, and chromatin accessibility analyses of *CHD1L* knock-out hiPSC-derived neuronal progenitor cells revealed that CHD1L regulates the expression levels and chromatin accessibility of genes involved in neuronal differentiation and synaptogenesis, including autism genes. Moreover, we found that *CHD1L* favors telencephalon development during forebrain regionalization by facilitating chromatin accessibility to pioneer transcription factors including SOX2 and OTX2 while simultaneously compacting chromatin through its interaction with the repressor NuRD complex. Last, atypical 1q21.1 CNV encompassing *CHD1L* and pathogenic missense and truncating *CHD1L* variants were found in individuals with autism. Overall, our data reveal a novel role for CHD1L as a master regulator of cell fate and its dosage imbalance contributes to the neuroanatomical and growth phenotypes associated with the 1q21.1 distal CNV.

**GRAPHICAL ABSTRACT:** 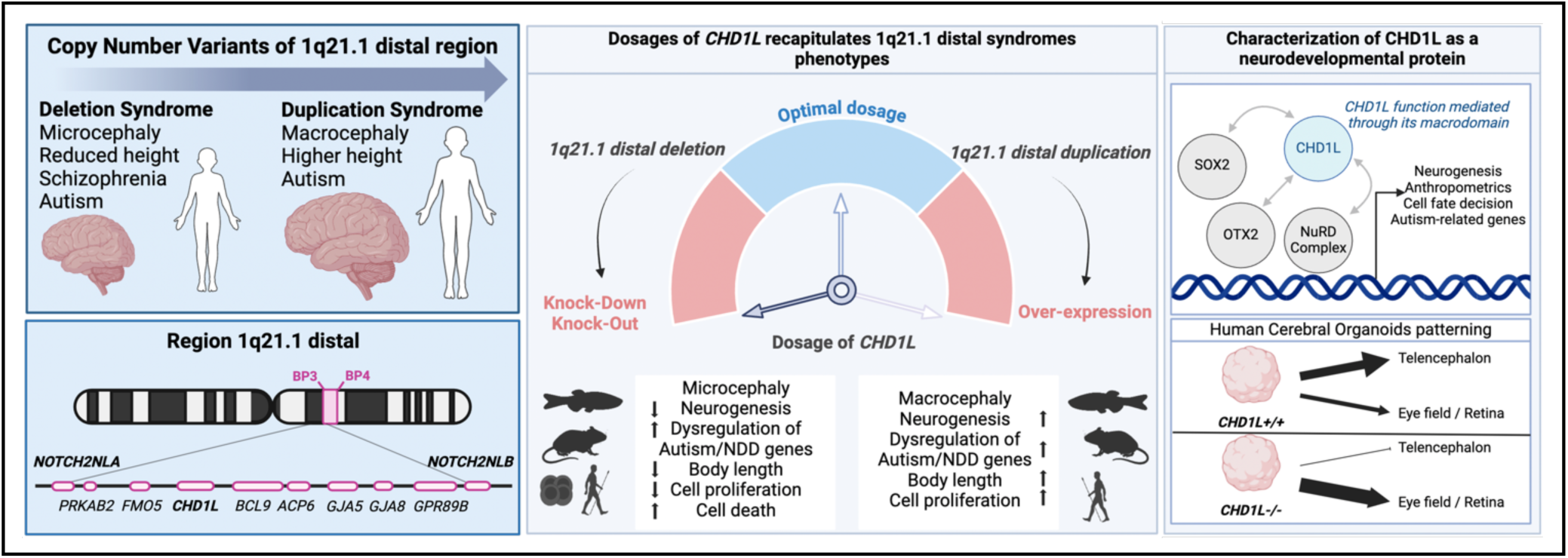

## INTRODUCTION

Copy number variants (CNV) are enriched in neurocognitive disorders, such as intellectual disability, schizophrenia, and autism spectrum disorders (ASD)(1–9), with a documented co-morbidity between CNV underscoring neurocognitive traits and micro/macrocephaly(10, 11). To date, a total of 17 autism-associated recurrent CNV loci are referenced (retrieved from the Simons Foundation Autism Research Initiative (SFARI); https://gene.sfari.org/database/cnv/). Among them, reciprocal deletions and duplications of the 1q21.1 breakpoint 3-4 (BP3-BP4) region are particularly prevalent in individuals with ASD.

The 1q21.1 region includes two CNV-prone regions, delimited by breakpoint BP2-BP3 and BP3-BP4 respectively, as well as deletion and duplication encompassing both intervals (categorized as class II rearrangements by Brunetti-Pierri in 2008)(4). Proximal BP2-BP3 microdeletions are known as susceptibility factor for thrombocytopenia-absent radius (TAR) syndrome (MIM: 274000) and brain anomalies have also been observed in some carriers^4^. The recurrent reciprocal 1q21.1 BP3-BP4 deletions (MIM: 612474) and duplications (MIM: 612475) also known as class I rearrangements or distal microdeletion/microduplication syndromes, have been found in individuals with syndromic autism (4, 12). Variable phenotypes have been reported, including congenital heart defects, autism, intellectual disability (ID), attention-deficit/hyperactivity disorder (ADHD), schizophrenia, increased/decreased head circumference and anthropometric variations(4). The 1q21.1 distal deletion is associated with microcephaly and short stature, whereas the reciprocal duplication is associated with increased risk of macrocephaly and carriers tend to be in the upper height percentiles(13, 14); suggesting a possible undergrowth/overgrowth phenotype. The 1q21.1 distal region contains eight protein-coding genes, including *PRKAB2, FMO5, CHD1L, BCL9, ACP6, GJA5, GJA8, GPR89B*. The human-specific *NOTCH2*-paralogs *NOTCH2NLA/B* provide the breakpoints of the 1q21.1 distal region. Among these genes, *GJA5* and *GJA8* encode gap junction proteins and missense variants at these loci have been reported to cause autosomal dominant atrial fibrillation and standstill (MIM: 614049 and 108770) and cataract (MIM: 11200) respectively(15–19).

A major challenge in the interpretation of CNV encompassing several genes is the identification of the loci whose dosage sensitivity drives the phenotype(s)(20). Only a handful of functional studies focusing on the 1q21.1 distal region syndromes have been reported(21, 22). First, with the advent of new technologies for generating cellular models from human differentiated cells, a recent study has indicated that human cortical neurons harboring the 1q21.1 distal deletion or duplication exhibit differentiation defects and altered electrophysiological property(21). Second, a mouse modeling 1q21.1 distal deletion is presenting schizophrenia-like behavior(22). Last, the human-specific *NOTCH2NLA and NOTCH2NLB* paralogs have been shown to control brain size *in vitro*(23, 24), however their candidacy as 1q21.1 phenotypic gene drivers remains unclear since no causative pathogenic single nucleotide variants have been reported for these genes in ASD cohorts.

Here, we aimed to identify the causal gene(s) that drive neuroanatomical and growth defects associated with the 1q21.1 distal CNV. To this end, we performed dosage perturbation of each of the 1q21.1 distal human gene during zebrafish development and identified *CHD1L* as a phenotypic driver. We then conducted transcriptomic, cistromic and chromatin accessibility analyses upon suppression of *CHD1L* in human induced pluripotent stem cell (hiPSC)-derived neuronal progenitor cells (hNPC), followed by cell fate assessment in human cerebral organoids and validation of pathogenicity of genetic variants in zebrafish.

CHD1L is a poly(ADP-ribose) and ATP-dependent remodeler with a key role in chromatin relaxation, and is implicated in several types of cancer(25–27). Structurally, it features a two-lobed catalytic Snf2- like ATPase domain linked to a C-terminal macrodomain *via* a linker region responsible for CHD1L binding to histones. CHD1L is widely recognized for its involvement in DNA-repair, particularly with PARP1 during Base Excision Repair (BER) and Nucleotide Excision Repair (NER) processes(28–35). Recent studies further clarified its role in early cellular response to DNA damage, revealing an auto- inhibitory mechanism arising from interactions between its macrodomain and the bilobate ATPase module(32, 36). In addition to its function in DNA repair, *Chd1l* has been shown to play a critical role in mouse embryo implantation(37) and to promote neuronal differentiation of cultured human embryonic stem cells(38). However, *CHD1L* has never been associated to neurodevelopmental syndromes. Here, we describe novel findings indicating that CHD1L plays a critical role as a co-transcription factor of key pioneer transcription factors to promote telencephalon fate in humans.

Overall, we found that dosage imbalance of *CHD1L* contributes to the 1q21.1 distal CNV-associated neuroanatomical and growth phenotypes. Furthermore, our study uncovers a hitherto unknown function of *CHD1L* that does not require its remodeling capability during human forebrain cell-fate decision, expanding its portfolio of functions beyond DNA repair and its association with cancer.

## MATERIAL AND METHODS

### Zebrafish strains and husbandry

Zebrafish (*Danio rerio*) were raised and maintained as described(39). Adult zebrafish were raised in 15 L tanks containing a maximum of 24 individuals, and under a 14 h to 10 h light-dark cycle. The water had a temperature of 28.5 °C and a conductivity of 200 µS and was continuously renewed. The fish were fed three times a day, with dry food and *Artemia salina* larvae. Embryos were raised in E3 medium, at 28.5 °C, under constant darkness. AB strain obtained from the European Zebrafish Resource Center (EZRC) was used as wild-type for this study, except for the *NOTCH2NLA/B* overexpression experiments for which AB strain obtained from the Zebrafish International Resource Center (ZIRC) was used as wild-type. Of note, a difference in baseline head size measurements between the two wild-type lines was observed (i.e. mean=120 µm for wild-type AB from EZRC and mean=140 µm for wild-type AB from ZIRC), which could potentially be explained by differences in genetic background. The mutant line *chd1l^sa14029^*, carrying a C>T point mutation at genomic location Chr 6: 36844273 (GRCz11) leading to a premature stop was obtained from ZIRC (ZIRC, ZL10486.01). F2 mutant embryos were obtained from an *in vitro* fertilization with heterozygous mutant F1 sperm (TL background) and AB eggs. F2 *chd1l^+/-^* fish were then inbred to generate F3 populations. All fish lines reproduce normally. Developmental stages of zebrafish embryos and larvae are indicated in the text and figures. For zebrafish embryos and larvae, both males and females were used since the sex can only be determined at 2 months of age. Genotyping of the *chd1l* mutants was performed by digestion with Taq alpha I restriction enzyme of the PCR product generated with the following primers: forward 5’- CAGCGTCAGTTTTGCTACCC - 3’; reverse 5’- CACCTGGATTGTTCTTGAGC - 3’. In figures, *chd1l*^+/-^ refers to heterozygous *chd1l ^sa14029^*^/+^ and *chd1l*^-/-^ refers to homozygous *chd1l^sa14029/sa14029^*. All animal experiments were carried out according to the guidelines of the Ethics Committee of IGBMC and ethical approval was obtained from the French Ministry of Higher Education and Research under the number APAFIS#15025-2018041616344504.

### *In vivo* analysis of gene expression and fish embryo manipulations

For overexpression experiments, the human wild-type mRNAs (*NOTCH2NLA*, ENST00000362074.8; *NOTCH2NLB*, ENST00000593495.4 ; *PRKAB2*, ENST00000254101.4; *CHD1L-202*, ENST00000369258.8, hereafter termed *CHD1L-FL*; *CHD1L-203*, ENST00000369259.4, hereafter termed *CHD1L-ΔLobe1*; *BLC9*, ENST00000234739.8; *ACP6*, ENST00000583509.7; *GJA5*, ENST00000579774.3; *GJA8*, ENST00000369235.1; *GPR89B*, ENST00000314163.12; *FMO5*, ENST00000254090.9) were cloned into the pCS2 vector and transcribed using the SP6 Message Machine kit (Ambion). Similarly, we cloned and transcribed two additional truncated forms of CHD1L, that were hereafter termed *CHD1L-FL-ΔMacro*, and *CHD1L-ΔLobe1-ΔMacro* using primers indicated in the STARMethods section. We injected 1 nl of diluted mRNA (50, 100 pg) into wild-type AB zebrafish embryos at the 1- to 2-cell stage. Injected embryos were fixed at 2 to 3 days post-fertilization (dpf) to perform whole-mount immunostaining or scored at 5 dpf for the head size and body length. The proxies that were used are the measurement of the distance between the eyes (distance between the convex tips of the eyecups) and the head size (distance from the anterior-most part of the forebrain until the hindbrain) on a dorsal view. The body length was measured from the head to the tail following the vertebral column and using inter-6 somites distance as a proxy. All the experiments were repeated at least three times and statistical analysis were performed as indicated to determine the significance of the observed phenotypes.

### High resolution episcopic microscopy and 3D reconstruction

5 dpf wild type AB and *chd1l^sa14029/sa14029^* homozygous larvae were fixed in Bouin’s solution, then washed and dehydrated with graded ethanol solutions (70%, 90%, 95%, 100%). The larvae were incubated in JB4 resin mixture (containing acridine orange and eosin)/ethanol (1:1), followed by incubation in pure resin mixture. The samples were embedded in fresh activated JB4 resin left to polymerize overnight at 4°C and harden in oven at 95°C. The blocks were sectioned using the Histo 3D system to generate data by repeated removal of 5 µm thick sections as described(40). Resulting HREM data with a voxel size of 4 X 4 X 5 µm^3^ were generated from approximately 250 aligned images. HREM data were analyzed using the Amira-Avizo™ software (Thermo Fisher Scientific) in segmentation mode to define brain volumes. The experiments were repeated twice, and statistical analysis were performed as indicated to determine the significance of the mutant phenotype.

### Whole-mount immunostaining on zebrafish larvae

2 dpf embryos were fixed in 4% PFA overnight and stored in 100% methanol at −20 °C. After rehydration in PBS, PFA-fixed embryos were washed in IF buffer (0.1% Tween-20, 1% BSA in PBS 1X) for 10 min at room temperature. The embryos were incubated in the blocking buffer (10% FBS, 1% BSA in PBS 1X) for 1 h at room temperature. After two washes in IF Buffer for 10 min each, embryos were incubated in the first antibody solution, 1:750 anti-phospho-histone H3 (ser10)-R, (Santa Cruz, sc-8656- R), in blocking solution, overnight at 4 °C. After two washes in IF Buffer for 10 min each, embryos were incubated in the secondary antibody solution, 1:1000 Alexa Fluor donkey anti-rabbit IgG, in blocking solution, for 1 h at room temperature. At least 15 larvae were imaged per condition and z-stacks were acquired. We used the ImageJ software to generate a "Maximum Intensity" projection and scored the number of fluorescent cells using the ITCN v1.6. plugin. The experiments were repeated three times, and statistical analysis were performed as indicated to determine the significance of the phenotypes.

### Whole-mount TUNEL assay on zebrafish larvae

TUNEL assay on 2 dpf embryos was performed using the ApopTag® Fluorescein In Situ Apoptosis Detection Kit (Merck Millipore, S7110) according to a modified protocol. Dechorionated embryos were fixed in 4% paraformaldehyde (PFA) at 4°C overnight and then stored in 100% methanol at −20°C for at least 2 hours. After rehydration in PBS, embryos were permeabilized with proteinase K (10µg/ml) in PBS for 5 minutes at room temperature and then washed 3 times in sterile water for 3 minutes each. Then, embryos were post-fixed with 4% PFA for 20 minutes at room temperature and followed by prechilled ethanol:acetic acid (2:1) for 10 minutes at -20°C. Embryos were washed in PBS-T (PBS 1X, 0.1% Tween-20) for 5 minutes, 3 times at room temperature. Incubation in equilibration buffer and next steps of the TUNEL were followed according to the manufacturer instructions. At least 15 larvae were imaged per condition and z-stacks were acquired. We used the ImageJ software to generate a "Maximum Intensity" projection and scored the number of fluorescent cells in defined regions of the zebrafish head (forebrain, midbrain, eyes) using the ITCN 1.6. plugin. The experiments were repeated three times, and statistical analysis were performed as indicated to determine the significance of the mutant phenotype.

### Acridine Orange whole-mount staining on zebrafish larvae

Acridine orange experiment was performed on 2 dpf embryos. The embryos were incubated at 28°C for 30 min in E3 embryo medium supplemented with 2 µg/mL Acridine Orange solution (Sigma, A231). After extensive washing, embryos were anesthetized with tricaine and imaged as Z-stacks with GFP green light excitation. Cell counting was performed with Fiji using the ITCN plugin coupled to manual counting. The experiments were repeated three times, and statistical analysis were performed as indicated to determine the significance of the mutant phenotype.

### Mouse *in utero* electroporation

*In utero* electroporation was performed using adult CD1 mice (Charles River) as previously described(41). Animal experimentations were performed at the IGBMC animal facilities in accordance with European Union and French legislation (APAFIS#17876-2018112916498297). The light/dark cycle for animals housed was 12h and all experiments were conducted during the light cycle. Timed pregnant mice (E13.5) were anesthetized with isoflurane (2 liters per min of oxygen, 4% isoflurane during sleep and 2% isoflurane during surgery operation; Tem Sega). The uterine horns were exposed, and a lateral ventricle of each embryo was injected using pulled glass capillaries (Phymep, 30-0016) and microinjector femtojet 4i (Eppendorf) with Fast Green (2 µg/ml; Sigma) combined with the following Endofree DNA plasmid solution: 1 μg of p-CIG-eGFP together with 1) 1 µg of pCAGGs-IRES-GFP empty or 2) 1 µg of wild-type mouse *Chd1l* cDNA (ENSMUST00000029730.4) cloned in pCAGGs-IRES-GFP or 3) 100 ng/µL of either control Rluc esi RNA (Sigma Aldrich, EHURLUC) or *CHD1L* targeting esi RNA (Sigma Aldrich, EMU070191). Plasmids were further electroporated into the neuronal progenitors adjacent to the ventricle by discharging five electric pulses at 30V for 50 ms at 950 ms intervals using an ECM 830 electroporator (BTX) and 3 mm electrodes. After electroporation, embryos were placed back in the abdominal cavity and development was allowed to continue until E14.5. Embryos brains were dissected and fixed in 4% PFA in PBS overnight. Brains were then rinsed in 20% sucrose to cryopreserve the tissues and embedded in sakura tissue-tek and stored at -80°C. Sectioning was performed using cryostat (Leica) at 14µm. Immunostainings were done using following antibodies concentrations: Stainings were performed using Anti-GFP (Abcam, ab6673, 1:500) + Donkey anti-goat 488 (Thermo Fisher, A-11055, 1:1000) and Anti-TBR1 (Abcam, ab31940, 1:250) + Donkey anti-rabbit 555 (Thermo Fisher, A-31572, 1:1000). Image acquisition was done with Leica confocal microscope SP5. Analysis was done by manual counting using Fiji. Electroporated and imaged embryos were obtained from at least three different litters, and statistical analysis were performed as indicated to determine the significance of the phenotypes.

### Cell culture, DNA transfection and single cell isolation by fluorescence-activated cell sorting (FACS)

Control human induced pluripotent stem cells (hiPSC GM8330-8, used in (*39*)), derived from adult fibroblasts, were kindly provided by Prof. M.E. Talkowski. The cells were maintained on Matrigel-coated dish (Corning) with mTESR™ (StemCell) and incubated at 37 °C in a humidified atmosphere with 5% CO2. We transfected the human iPSCs with the pSpCas9(BB)-2A-GFP gRNA plasmid using Lipofectamine™ Stem Reagent, adapting the protocol described in(42). At 48 hours post-transfection, the hiPSCs were dissociated into a single cell suspension with Accutase and resuspended in PBS with 10 μM ROCK inhibitor (Santa Cruz). All samples were filtered through 5mL polystyrene tubes with 35 µm mesh cell strainer caps (BD Falcon, 352235) immediately before being sorted. After adding the viability dye DAPI (BD Bioscience), single GFP+ DAPI- cells were isolated by fluorescence-activated cell sorting (FACS) gated for a high level of GFP expression and sorted, with one cell placed into each well of Matrigel-coated 96-well plates by BD FACS Aria II with 100-mm nozzle under sterile conditions. The medium was supplemented with CloneR™ (StemCell) from Day 0 to Day 4 according to the manufacturer’s instructions.

### hiPSC-editing guide RNA design and preparation

We used the CRISPR MIT tool (http://crispr.mit.edu) to generate a guide RNA targeting the exon 1 of *CHD1L* (5’-TCATACTGAGGGCCGAGCCGAGG-3’, chr1: 147242763-147242785, GRCh38). The gRNA was cloned into pSpCas9(BB)-2A-GFP (Addgene, PX458) plasmid. Validation of the guide sequence in the gRNA vector was confirmed by Sanger Sequencing. Before transfection, all plasmids were purified from PureLink™ HiPure Plasmid Midiprep Kit according to the manufacturer’s instruction (Thermo Fisher Scientific).

### Colony screening and western blot validation

Genomic DNA from two thirds of each hiPSC colony (obtained ∼14 days after sorting) were extracted by using Quick-DNA 96 kit (Zymo research) and screened by PCR (using the indicated primers: forward 5’- GGAAGTTGGGAGGGAGGT-3’ and reverse 5’ GCTGATCTCACCACGTTTCC-3’) followed by Sanger sequencing. For hiPSC screening validation, 100 μg of total iPSC protein lysate was prepared in RIPA buffer and Protease Inhibitor Cocktail from control and two *CHD1L*-edited iPSCs lines, diluted 1× final with Laemmli buffer and DTT 0.1 M, boiled for 5 min and then separated by SDS-PAGE on 10% polyacrylamide gels. Resolved proteins were transferred to nitrocellulose membranes and blocked in 3% milk 1× TBS for 1 h at room temperature prior to incubation with either anti-CHD1L (2170C3a) antibody (Santa Cruz, sc-81065, 1:200) or anti-β-Tubulin (1:10,000, produced in house). Membranes were washed and incubated in goat anti-mouse peroxidase secondary antibody (Jackson Immuno Research, 1:10,000). Blots were developed using Immobilion Western (Millipore, France) according to the manufacturer’s instructions.

### Karyotyping

Karyotypic analysis was performed by Cell Guidance Systems (Cambridge, UK) following standard methods.

### Differentiation of hiPSCs into Neural Progenitor Cells (hNPC)

Cells were differentiated into hNPC utilizing the StemXVivo Neural Progenitor Differentiation Kit (R&D systems). The expression of CNS-type hNPCs markers was validated by immunocytochemistry, as described in the Fixing and Staining procedure of the StemXVivo datasheet, using anti-Human SOX1 included in the kit and Sheep Anti-Human Pax6 Polyclonal Antibody (R&D Systems, AF8150) (10 µg/ml in PBS) followed by incubation with Chicken anti-Goat IgG (H+L) Cross-Adsorbed Secondary Antibody, Alexa Fluor 488 and Donkey anti-Sheep IgG (H+L) Cross-Adsorbed Secondary Antibody, Alexa Fluor 594 (Thermo Fisher Scientific, 1:200), respectively.

### RNA-seq library preparation and bioinformatic analyses

Total RNA was extracted using RNA Plus Mini Kit (Macherey Nagel). Libraries were prepared using the Illumina TruSeq stranded mRNA protocol and sequenced on an Illumina HiSeq 4000 (single-end 50 bp reads). Reads were preprocessed using cutadapt (v1.10) (43) in order to remove adapter, polyA and low-quality sequences (Phred quality score below 20) and reads shorter than 40 bases were discarded for further analysis. Reads mapping to rRNA were also discarded. This mapping was performed using bowtie (v2.2.8) (44). Reads were then mapped onto the hg38 assembly of Homo sapiens genome using STAR (v2.5.3a) (45). Gene expression quantification was performed using htseq-count (v0.6.1p1) (46), with annotations from Ensembl version 98. Normalization of read counts and differential expression analysis between controls and CHD1L-edited samples were performed using the method proposed by Love et al. (47) and implemented in the Bioconductor package DESeq2 (v1.16.1), adjusting for batch effect. For comparison among datasets, transcripts with more than 50 raw reads were considered. The common differentially expressed genes (DEG) discussed in this manuscript are those that show differential expression in both CHD1L mutant lines compared to the control line, and exhibit the same direction of change (either up-regulated or down-regulated) in both mutants. For STRING analysis, the online tool was used (string-db.org). The database DECIPHER was also used for gene annotations (decipher.sanger.ac.uk/ddd/ddgenes). The top 500 DEG (both upregulated and downregulated and filtered based on FDR) were entered into Online tool Lisa.cistrome for Transcription Regulators prediction (lisa.cistrome.org). IGV v2.17.4 was used to generate sashimi plot. For Gene Ontology terms analysis, the online tool PANTHER was used (geneontology.org), and illustrated using R package ggplot2.

### ATAC-seq library preparation

Assay for transposase accessible chromatin was performed on hiPSC-derived hNPC using a derived protocol from Buenostro et al. (48). A total of 50,000 cells from GM8330-8 (control line *CHD1L^+/+^*), Line 1 and Line 2 (isogenic *CHD1L* mutant lines) lines were resuspended in resuspension buffer (10mM Tris-HCl pH 7.5, 10mM NaCl, 3mM MgCl2) and then lysed in lysis buffer (Resuspension buffer + 0.1% NP40 + 0.1% tween-20+ 0.01% Digitonin) and incubated 3 min on ice. Then, 1 mL of wash buffer was added (resuspension buffer + 0.1% Tween-20). Cells were centrifugated 10 min at 500G at 4°C and supernatant was discarded. Transposition reaction mix (Illumina, Tagment DNA Enzyme and Buffer Small Kit, 20034197) was added to pellet and incubated 30 min at 37°C in a thermomixer at 1,000 rpm. DNA fragments were isolated using Qiagen MinElut Reaction Cleanup kit. DNA fragments were amplificated and libraries were generated by PCR using appropriate primers as described in^54,55^ and NEBNext High-Fidelity 2X PCR Master mix (NEB, M0541S). Finally, libraries were purified using SRIselect beads (Beckman Coulter) by a one-sided purification to remove primers. Sequencing was performed using Illumina HiSeq 4000 with 100 bp paired-end sequencing.

### ATAC-seq bioinformatic analyses

Data analysis was performed using the Encode ATAC-seq pipeline (v1.4.2). Adapter sequences were removed and low-quality ends were trimmed. Reads were mapped onto the hg38 assembly of Homo Sapiens genome using Bowtie2 (v2.2.6) (44) choosing the zero multi-mapping option. Mitochondrial reads were removed. The Peak calling was performed using MACS2 (v2.1.1.20160309). Finally, the optimal overlap peaks were used for downstream analyses. Peaks from different conditions were merged to form a consensus peak set. The peaks were annotated using annotatePeaks.pl script in Homer program (49) and with Ensembl 98 database. The read coverage for each sample was calculated with multicov function from bedtools program (v2.26.0). Differential analyses of Control GM8330-8 vs either Line 1 or Line 2 isogenic mutants were performed using the Bioconductor package DESeq2 (v1.16.1) (47). The common differentially accessible peaks discussed in this manuscript are those that show differential accessibility in both CHD1L mutant lines compared to the control line, and exhibit the same direction of change (either more accessible or less accessible) in both mutants. For the TOBIAS analysis (Transcription factor occupancy prediction by investigation of ATAC-seq signal) of enriched motif elements, the pipeline snakemake (v0.12.11) was used(50). ATACseq datasets were compared to H3K4me2 peaks from CUT&RUN analysis performed in this study using seqMINER(51). Representative traces were generated using Figeno(52).

### CUT&RUN library preparation

Protocol was adapted from Hainer and Fazzio (53). For each condition, a total of 1.10^5^ hNPC cells were washed with in cold PBS 1X and resuspended in cold Nuclear Extraction Buffer (20mM HEPES-KOH, pH 7.9; 10mM KCl; 0.5mM Spermidine; 0.1% TritonX-100; 20% glycerol; PIC 1X) and incubated for 5 min at 4°C. The nuclei obtained were pelleted and recovered in cold NEB. Concanavalin A magnetic beads (BioMagPlus, 86057-3, 25 μL bead slurry/sample) were washed twice in cold Binding Buffer (20mM HEPES-KOH, pH 7.9; 10mM KCl; 1mM CaCl2; 1mM MnCl2; 1X PIC) and recovered in Binding Buffer. The beads were added to the nuclei with a gentle vortexing and incubated for 10 min on the wheel at 4°C. Bead-bound nuclei were incubated for 5 min at room temperature in Blocking Buffer (1mL Wash Buffer (20mM HEPES-KOH, pH 7.5; 150mM NaCl; 0.5mM Spermidine; 0.1% BSA; PIC 1X), 4 μL 0.5M EDTA) and washed in Wash Buffer. Nuclei were re-suspended in 250 μL of cold Wash Buffer per condition. An antibody solution (250 µL) containing the primary antibodies anti-H3K4me2 or anti-CHD1L diluted at 1:100 was added to the bead-bound nuclei with a gentle vortexing. Samples were incubated overnight at 4°C on a wheel. The beads were then washed with cold Wash Buffer and protein A–micrococcal nuclease recombinant protein (pA–MN) enzyme (0.7ng/ml; 200 µL per condition) was added by gentle vortexing. Samples were incubated with rotation at 4 °C for 1 h. The pA–MN was produced in-house according to the protocol described by Schmid et al. (54) and using the pK19pA–MN plasmid (RRID: Addgene_86973; http://n2t.net/addgene:86973). The nuclei were then washed and re-suspended in Wash Buffer. The samples were incubated for 10 min in equilibrated water at 0°C. Cleavage was initiated by the addition of 100mM CaCl2 with gentle vortexing and incubated for 30min. The reaction was stopped by adding 2X Stop Buffer (5M NaCl; 0.5M EDTA; 0.2M EGTA; 50 μg/μL RNAseA; 40 μg/mL glycogen) and DNA fragments were released by passive diffusion during incubation at 37 °C for 20 min. After centrifugation for 5 min at 16,000g at +4 °C to pellet cells and beads, 3 μL 10% SDS and 2.5 μL Proteinase K 20 mg/mL were added to the supernatants, and samples were incubated 10 min at 70 °C. DNA purification was done with phenol/chloroform/isoamyl alcohol extraction followed by a second chloroform extraction. DNA was precipitated with ethanol after addition of 20 μg glycogene and resuspended in 0.1X Tris EDTA. DNA fragments were amplificated and library was generated by PCR using MicroPlex Library Preparation Kit v3 (Diagenode, C05010001). Finally, libraries were purified using SRIselect beads (Beckman Coulter) by a one-sided purification to remove primers. Sequencing was performed using NextSeq 2000 with 50 bp paired-end sequencing.

### CUT&RUN bioinformatic analyses

Paired-end reads were mapped to Homo Sapiens genome (assembly hg38) using Bowtie2 (v2.3.4.3, parameters: -N 1 -X 1000). Reads overlapping with ENCODE hg38 blacklisted region V2 were removed using bedtools. Bigwig tracks were generated using bamCoverage. Tracks were normalized with RPKM method. The ≤120 bp fragments were used for samples obtained with anti-CHD1L. Peak calling was performed with the Sparse Enrichment Analysis for CUT&RUN (SEACR v1.3) tool using a numeric threshold of 0.002 for CHD1L (https://seacr.fredhutch.org), and with MACS2 for H3K4me2 (broad peaks, q-value 0.05, other parameters as default). Peak annotation and genomic features (promoter/TSS, 5’ UTR, exon, intron, 3’ UTR, TTS and intergenic regions) were defined and calculated using Refseq and HOMER according to the distance to the nearest TSS. HOMER and RSAT were used for motif search(55). Heatmaps were generated with deeptools(56). ChIP-Atlas was used for Transcription Factor enrichment (chip-atlas.org)(57) using the following parameters: ChIP TFs and others for Experiment type, Neural for Cell type Class, 50 for Threshold for significance, and random permutation. CHD1L CUT&RUN peaks were compared to the following ChIP-seq data: SOX2 peaks from human neural progenitor cells (SRX330107), CHD7 peaks from human cerebellum (SRX9795022), HDAC2 (SRX19212560) and RBBP4 (SRX19212562) peaks from human neuroblastoma. Representative peaks were generated using Figeno(52) on the reference human genome *hg38*. Intersections between datasets were performed with Venny 2.1 (https://bioinfogp.cnb.csic.es/tools/venny/). WebGestalt was used to run DisGeNET enrichement analysis for *CHD1L* and *SOX2* common gene targets (WebGestalt.org).

### Immunoprecipitation and LC-MS/MS

Control *CHD1L^+/+^* and mutant *CHD1L^-/-^* (Line 1) hNPCs were harvested in PBS and centrifugated at 300g for 5 min. Pellets were lysed for 30 min under agitation in lysis buffer (Tris pH7.5 50mM, NaCl 150mM, Glycerol 5%, NP40 1%, PIC 1X in H20). Immunoprecipitation were performed using Slurry Dynabeads G coupled to 5 µg of Anti-CHD1L antibody for 1h at 4°C under agitation. On ice, 100 μg of proteins were added on antibody-coupled beads and agitated at 4°C overnight. After three washes in lysis buffer, bound proteins were eluted in 1X blue Laemmli and 0.1M DTT and boiled. Experiment was repeated three times for each genotype. The isolated proteins were subjected to an overnight trypsin digestion at 37°C. The peptides were separated on a C18 column using an Ultimate 3000 nano-RLSC (Thermo Fisher Scientific) coupled to an LTQ-Orbitrap Elite mass spectrometer (Thermo Fisher Scientific) for peptide identification. Proteins were identified with Proteome Discoverer 2.4 software (Thermo Fisher Scientific, PD2.4(58)) from Homo Sapiens proteome database (Swissprot, reviewed, release 2021_06_03, 20309 entries), and proteins were quantified with a minimum of 2 unique peptides based on the XIC (sum of the Extracted Ion Chromatogram). Partners of CHD1L were considered if enriched (Fold-Change >1) in control condition compared to the protein listed in the mutant condition to exclude nonspecific bindings. Protein identified were filtered for nuclear protein only using BioDBnet.org(59) db2db tool. For visual representation and Gene Ontology terms analysis, the online tool STRING.db(60) was used, and illustrated using R packages GGplot2.

### Co-immunoprecipitation on HEK cells

*CHD1L* and *SOX2* cDNA were cloned in pCDNA3.1 plasmids with either N-terminal HA or FLAG tag sequence respectively. Plasmids coding for *HA-CHD1L* or *FLAG-SOX2* were transfected in HEK293-T using lipofectamine 2000 (0.5 µg alone or 1.5 µg when co-expressed). The next day, cells were lysed in Lysis Buffer (RIPA, 50mM Tris-HCL pH 8, 150mM NaCl, 0.5% TritonX-100 and 1X PIC). Samples were sonicated and centrifuged for protein purification. Proteins were incubated with anti-HA magnetic beads overnight at 4°C with agitation. After washes, bound proteins were diluted 1× final with Laemmli buffer and DTT 0.1 M, boiled for 10 min and then separated by SDS-PAGE. Resolved proteins were transferred to nitrocellulose membranes and blocked in 5% milk 1× TBS-Tween 0.1% for 1 h at room temperature prior to incubation with rabbit anti-HA and rat anti-FLAG antibodies. Membranes were washed and incubated in goat anti-rabbit and Goat anti-rat peroxidase secondary antibody (1:10000, Jackson Immuno Research, U.K.). Blots were developed using Immobilon Western (Millipore, France) according to the manufacturer’s instructions. Experiment was performed three times.

### Generation of human cerebral organoids and immunostaining

Human cerebral organoids (hCO) were generated using the STEMdiff Cerebral Organoid Kit (STEMCELL, 08570) and following the manufacturer’s instructions, and matured until Day 60 *in vitro*. For immunostainings, cerebral organoids were fixed in 4% PFA overnight at 4°C, after PBS washes, cerebral organoids were incubated in PBS-sucrose 30% until sedimentation. The hCO were embedded in 1:1 OCT and Sucrose 30% and stored at -80°C. Cryosections were performed at 14 µm. Sections were hydrated in PSB 1X for 10 min then permeabilized with 0.1% Triton in PBS for 20 min at RT and were incubated 1 hour with Blocking Solution (10% FCS + PBS-Triton 0.3%). Sections were incubated overnight at 4°C with primary antibodies anti-SOX2 (Santa Cruz, sc-365823, 1:200) and anti-TUJ1 (D71G9) (Cell Signaling technology, 5568S, 1:500) in blocking solution. After extensive washing, sections were incubated in blocking solution with secondary antibodies anti-mouse 488 (Thermo Fisher, A1101, 1:500) and anti-rabbit 647 (Thermo Fisher, A21244, 1:500). Images were acquired using Leica Image acquisition Leica confocal microscope SP8X. Image reconstruction was done using FiJi.

### Western Blot for CHD1L and TUJ1 in cerebral organoids

Human cerebral organoids were snap frozen using liquid N2 and lysed in 200 µl of RIPA buffer supplemented with Protease Inhibitor Cocktail. Proteins were purified and quantified, then resuspended in 1× final with Laemmli buffer and DTT 0.1 M, boiled for 5 min and 15 µg of proteins was separated by SDS-PAGE on 10% polyacrylamide gels. Resolved proteins were transferred to nitrocellulose membranes and blocked in 3% milk 1× TBS for 1h at room temperature prior to incubation with either anti-CHD1L (2170C3a) antibody (Santa Cruz, sc-81065, 1:200), anti-β tubulin (1:10,000, produced in house) and anti-TUJ1 (D71G9) (Cell Signaling technology, 5568S, 1:1000). Membranes were washed and incubated in goat anti-mouse peroxidase secondary antibody (Jackson Immuno Research, 1:10,000) or goat anti-rabbit (for TUJ1) peroxidase secondary antibody (Jackson Immuno Research, 1:10,000). Blots were developed using Immobilon Western (Millipore, France) according to the manufacturer’s instructions.

### Single-nuclei Multiome on human cerebral organoids

Human cerebral organoids were maintained until day 60 *in vitro*. Nuclei isolation was followed the protocol “Nuclei Isolation from Complex Tissues for Single Cell Multiome ATAC + Gene Expression Sequencing” (10Xgenomics, protocol CG000375 Rev C) without the cell sorting step. Specifically, a total of six cerebral organoids per condition, control *CHD1L*^+/+^ and *CHD1L*^-/-^ (Line 1), were washed in PBS 1X. Lysis were performed using 1X NP40 Lysis Buffer with mechanical dissociation of tissues. Samples were filtered through 70 µm strainer and centrifugated for 5 min at 4°C 500 rcf. Pellets were re-suspended in PBS + 1% BSA + 1U/µl RNAse Inhibitor and incubated for 5 min. After a second centrifugation, pellet was re-suspended in 0.1X Lysis Buffer and incubated 5 min on ice. Wash buffer was added, and samples were centrifugated before a final resuspension in Diluted Nuclei Buffer. Libraries were generated following the 10X Genomics Multiome Guidelines (protocol vA.01) and the protocol “Chromium Next Gem Single Cell Multiome ATAC + Gene expression User guide” (CG000338 Rev E). Sequencing was performed using Illumina NextSeq 2000 with 50 bp paired-end sequencing for ATAC-seq and 28 + 85 bp paired-end sequencing for RNA-seq. BCLconvert (v3.8.4), Cell Ranger ARC (v2.0.2) and the human reference cellranger-arc-GRCh38-2020-A-2.0.0 were used for demultiplexing, alignment, barcode and UMI filtering and counting. Control *CHD1L^+/+^* and *CHD1L*^-/-^ samples were aggregated using cellranger-arc aggr function without performing the depth normalization.

### Seurat analysis of snMultiome

10x Genomics multiomic data were processed using Seurat (v4.3) and Signac (v1.1) (61). The output of the Cell Ranger arc pipeline was read using the Read10X function of Seurat (R, v4.2.2) to obtain a matrix of the number of UMIs of each gene detected in each cell. A total of 4,019 cells, including 2,385 cells from *CHD1L*^+/+^ organoids and 1,634 cells from *CHD1L*^-/-^ organoids, were retained after filtering based on RNA-assay metrics (total counts > 1,000 and < 50,000, percent.mt < 20) and ATAC-assay metrics (total counts > 2,000 and < 50,0000). We applied SCTransform to normalize RNA counts and TFIDF to normalize ATAC peaks. Dimensionality reduction was performed using Latent Semantic Indexing (LSI) for ATAC data and Principal Component Analysis (PCA) for RNA data. We constructed a weighted nearest neighbor graph (WNN) with Seurat (v4.0)(62) using 2-50 ATAC LSI dimensions and 1-50 RNA PCA dimensions. We used the FindClusters function with the SLM algorithm and a resolution of 0.8 to identify the clusters. Differentially expressed genes (DEG) for RNA-seq analysis were calculated using the FindAllMarkers function with the Wilcoxon test. For the Sankey diagram, A threshold of read number > 1 was used to count the number of positive nuclei expressing *FOXG1* and *SIX3*. Representative figures for genotype or marker gene expressions were generated using R and ggplot2.

### SCENIC+ analysis of snMultiome

Multiomic datasets were subjected to the SCENIC+ workflow (v0.1.dev456+g9662363) as described by Bravo Gonzalez-Blas et al., 2023(63). Topic modeling of scATAC-seq data was carried out using pycisTopic (v1.0.2.dev15+g242c2a4). Consensus peaks were called using Macs2 (total of 362,610 peaks), and topic modeling was performed using Latent Dirichlet Allocation (LDA) with the collapsed Gibbs sampler. A model of 16 topics was selected based on stabilization of metrics. Motif enrichment analysis was conducted using pycisTarget (v1.0.2.dev11+g7daf370). The hg38 v10 motif collection was downloaded from the cistarget resources website (https://resources.aertslab.org/cistarget/). Motif enrichment was performed using both the cisTarget and DEM algorithms on cell line-based DARs with default thresholds. The analysis was run both including and excluding promoters, defined as regions within 500 bp up- or downstream of the TSS of each gene. The SCENIC+ workflow was executed using default parameters. A search space of a maximum between either the boundary of the closest gene or 150 kb and a minimum of 1 kb upstream of the TSS (ensemble release 98) or downstream of the end of the gene was considered for calculating region–gene relationships using gradient-boosting machine regression. TF–gene relationships were calculated using gradient-boosting machine regression between all TFs and all genes. Genes were considered as TFs if they were included in the TF list available on http://humantfs.ccbr.utoronto.ca/ (v1.01) (64). Final eRegulons were constructed using the GSEA approach, and only eRegulons with a minimum of ten target genes were retained. For each eRegulon, cellular enrichment scores (AUC) of target genes and regions were calculated using the AUCell algorithm (65). eRegulons with correlation coefficients between pseudobulked per cell type TF expression and region enrichment AUC scores >0.6 or <-0.4 were considered high quality and used for downstream analysis. This resulted in 63 regulons, with a median of 408 genes and 719 regions per regulon. The eRegulon enrichment scores for regions and genes were normalized for each cell and used as input into t-distributed stochastic neighbor embedding (*t*-SNE) from the Python package fitsne (v.1.2.1). eRegulon specificity scores were calculated, per cell type and eRegulon, using the RSS algorithm as described in (66), using target region or target gene eRegulon enrichment scores as input.

### Patient recruitment

Recruitment of the affected individual to this study was initiated by clinicians, in the context of care, in accordance with the Montpellier Hospital ethics committee, followed by written informed consent from his legal representatives. The patient genetic variants were determined with exome sequencing followed by CGH array. The exome sequencing was performed with a NextSeq 550 sequencer (Illumina) in paired end 2x75 base-pairs. A bioinformatics filter based on the individual’s symptoms is applied to prioritize already known genes or those expressed in the brain. Three heterozygous variants were identified in the individual: NM_004284.6 (*CHD1L*): c.1929del, p.Arg643Serfs*16; NM_001346810.2 (*DLGAP2*): c.1696C>T, p.Arg566*; NM_017990.5 (*PDPR*):c.1147G>T, p.Gly383*.

### SDS-Page analysis of p.Arg392His CHD1L

The cDNA coding for the various constructs of *CHD1L* wild-type and *CHD1L* carrying the p.Arg392His mutation were cloned into a pET-MCN vector(67) coding for an N-terminal 6xhistidine affinity purification tag. *E. coli* BL21(DE3) cells were transformed with the respective plasmids and the colonies having incorporated the plasmids were selected by ampicillin resistance. These colonies were used for inoculating 4 mL liquid cultures. The cultures were left to grow upon shaking during 4h at 37°C. The cultures were then cooled down at 20°C and protein expression was induced by adding 0.5 mM final of IPTG (Euromedex). Expression was achieved overnight upon shaking at 20°C. The cultures were centrifuged (2,500 rpm, 20 min) and the supernatant discarded. All cultures were resuspended in a purification buffer composed of 10 mM Tris pH 8.0 and 200 mM NaCl. A small sample of each resuspended cultures was collected, mixed with Laemmli buffer and boiled at 95°C for 5 min prior to analysis of the total expression levels by SDS-PAGE. The resuspended cultures were lysed by sonication. After centrifugation (2,500 rpm, 20 min), the supernatants were incubated with 20 µL of Talon affinity beads (Clonetech) for 1h at 4°C. The supernatants were then discarded, and the beads washed twice with the purification buffer. The beads were then resuspended in Laemmli buffer for analysis of the soluble levels by SDS-PAGE.

### Statistical analysis

Number of measures (n), statistical tests used, and associated *p*-values are indicated in the figures or in the legends of each figure. Graph representation and statistical analyses were performed using GraphPad Prism 10 (v10.2.0). All measurements were performed on distinct samples. Rout Test was performed to remove outliers. Graphs are represented as Mean ± Standard Error of the Mean (SEM). Statistical tests are mentioned in figure legends. To ease the visualization of the data, the non-significant *p*-values are not showed on the graphs.

## RESULTS

### *In vivo* testing of the human 1q21.1 distal region genes

Manipulation of zebrafish embryos is an attractive alternative method to discover human dosage-sensitive genes. This is particularly useful when the CNV under investigation has mirrored anatomical phenotypes detectable during early development, allowing for assays using a combination of gene suppression and overexpression experiments(68–70). Given the association between the 1q21.1 distal CNV and changes in head circumference and anthropometrics traits, we proposed that *i*) systematic overexpression of each of the ten genes comprised in the region might yield a set of transcripts causing macrocephaly as described in 1q21.1 distal duplication syndrome and *ii*) reciprocal suppression of these candidate genes should yield the microcephalic phenotype seen in the 1q21.1 distal deletion syndrome.

To test this possibility, we first queried the zebrafish genome by reciprocal BLAST (Basic Local Alignment Search Tool) for each of the eight single copy protein coding genes within the 1q21.1 distal region (**Figure 1A**). All genes but *FMO5* have Ensembl-annotated orthologues (CABZ01083448.1/*prkab2* (ENSDARG00000067817); *chd1l* (ENSDARG00000015471); *bcl9* (ENSDARG00000036687); *gpr89b* (ENSDARG00000077983); *acp6* (ENSDARG00000040064)); for two of them, *GJA5* and *GJA8*, two copies each were mapped (*gja5a* (ENSDARG00000040065), *gja5b* (ENSDARG00000069450) and *gja8a* (ENSDARG00000069451), *gja8b* (ENSDARG00000015076), respectively). *NOTCH2*-derived *NOTCH2NLA* and *NOTCH2NLB* paralogs, located at the BP3-BP4 breakpoints of the 1q21.1 distal region, are human specific genes(23, 24).

**Figure 1:**
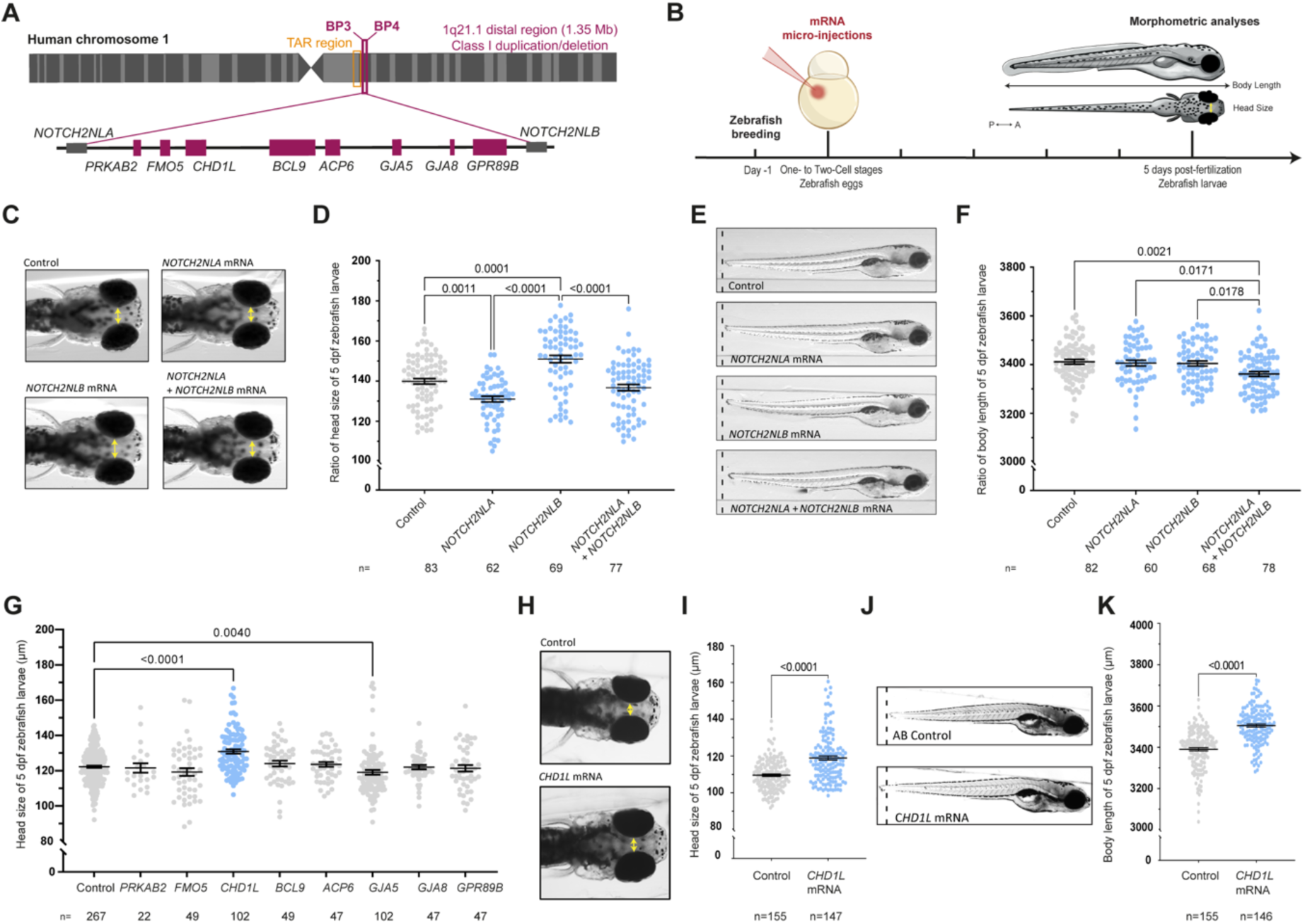
*In vivo* overexpression of the genes from the 1q21.1 distal region in zebrafish. **(A)** Schematic representation of the human 1q21.1 BP3-BP4 distal region showing eight genes comprised in the region and *NOTCH2*-paralogs *NOTCH2NLA* and *NOTCH2NLB* located at the breakpoints. **(B)** Schematic representation of the experimental procedure including micro-injection of mRNA in one- to two-cell stage zebrafish eggs and morphometric analyses at 5 days post-fertilization (dpf) for the head size and the body length of zebrafish larvae. **(C)** Dorsal views of the head of control larvae and larvae injected with *NOTCH2NLA* mRNA (50pg), *NOTCH2NLB* mRNA (50pg) or *NOTCH2NLA*+*NOTCH2NLB* mRNA (50+50pg) at 5 dpf. Double-ended arrows indicate distance between the eyes. **(D)** Dot plot showing the distance between the eyes (head size) of control larvae and larvae injected with *NOTCH2NLA* mRNA (50pg), *NOTCH2NLB* mRNA (50pg) or *NOTCH2NLA*+*NOTCH2NLB* mRNA (50+50pg) at 5 dpf. Data shown as mean ± SEM of a least triplicate batches; Kruskal-Wallis test. **(E)** Lateral view of control larvae and larvae injected with *NOTCH2NLA* mRNA (50pg), *NOTCH2NLB* mRNA (50pg) or *NOTCH2NLA*+*NOTCH2NLB* mRNA (50+50pg) at 5 dpf. **(F)** Dot plot showing the body length measurements of control larvae and larvae injected with *NOTCH2NLA* mRNA (50pg), *NOTCH2NLB* mRNA (50pg) or *NOTCH2NLA*+*NOTCH2NLB* mRNA (50+50pg) at 5 dpf. Data shown as mean ± SEM of at least triplicate batches per condition; Ordinary One-way ANOVA test. **(G)** Dot plot showing the distance between the eyes (head size) of 5 dpf control larvae and larvae injected with each of the 8 human genes from the 1q21.1 distal region. Data shown as mean ± SEM of a least triplicate batches per gene; Kruskal-Wallis test. **(H)** Dorsal views of control and larvae injected with *CHD1L* mRNA (100pg) at 5 dpf. Double-ended arrows indicate distance between the eyes. **(I)** Dot plot showing the distance between the eyes (head size) of control and *CHD1L* mRNA-injected larvae (100pg) at 5 dpf. Data shown as mean ± SEM of a least triplicate batches; Mann-Whitney test. **(J)** Lateral views of control and *CHD1L* mRNA-injected larvae (100pg) at 5 dpf. **(K)** Dot plot showing the body length measurements of control or injected larvae with *CHD1L* mRNA (100pg) at 5 dpf. Data shown as mean ± SEM of triplicate batches; Mann-Whitney test. To ease the visualization of the data, the non-significant *p*-values are not showed on the graphs.

Using a similar strategy to our previous studies (68–70), we first generated capped messenger RNA (mRNA) for all 10 human genes and injected zebrafish embryos at one-cell stage with a dose of 50 pg or 100 pg of mRNA per egg. To achieve expression above the baseline of any single transcript, we typically used an injection amount corresponding to 0.25%–0.5% of the total poly(A) mRNA found in a zebrafish embryo. Previous study indicated that the injected human mRNAs are persisting in the larvae up to 4.25 days post-fertilization (dpf)(68) (**Figure 1B**).

First, we determined whether *NOTCH2NLA* and *NOTCH2NLB* overexpression might affect brain size and/or body length of zebrafish larvae. Despite the absence of *NOTCH2NLA/B* in zebrafish, their overexpression led to relevant phenotypes. More specifically, the overexpression of each of the two genes led to opposite effects on head size: *NOTCH2NLA* overexpression led to microcephaly (-6.3%) whereas *NOTCH2NLB* overexpression led to macrocephaly (+7.9%) in overexpressant larvae compared to controls (**Figure 1C and D**). Strikingly, overexpression of both *NOTCH2NLA* and *NOTCH2NLB* did not lead to head size variations, suggesting a compensatory effect (**Figure 1C and D**). In addition, we observed no physiologically relevant effect on the body length upon injection of *NOTCH2NLA*, *NOTCH2NLB*, or both transcripts (-1.5%) in zebrafish (**Figure 1E and F**). Despite their critical role during primate brain development (23), our *in vivo* data indicated that the opposite effects mediated by the two *NOTCH2NL* paralogs counteract each other under control conditions. Moreover, we exclude a role of these genes for the growth phenotypes observed in individuals with 1q21.1 distal syndromes. Therefore, we postulated that other genes comprised in the 1q21.1 distal region contribute to the 1q21.1 distal syndromes head size and growth phenotypes.

We thus overexpressed each of the 8 genes comprised in the 1q21.1 distal region in zebrafish. We did not observe toxicity, lethality, or gross morphological defects upon injection, except for *GJA5*, for which the maximum dosage that could be tested was 50 pg. *GJA5* is a member of the connexin gene family, whose mutations have been associated with atrial fibrillation and arteriovenous malformations. The gene is likely responsible for the congenital heart diseases associated with the 1q21.1 distal deletion, whereas in case of a duplication of *GJA5,* tetralogy of Fallot is more common(15–17). Strikingly, only the overexpression of the chromatin remodeler *CHD1L* yielded a significant increase of the head size at 5 dpf (**Figure 1G**). Specifically, we measured the distance between the eyes (**Figure 1H and I**) and the distance between the anterior-most part of the forebrain and the hindbrain (**Supplementary Figure S1A and B**) in *CHD1L*-injected larvae, revealing a brain overgrowth phenotype when *CHD1L* is overexpressed. We further evaluated whether *CHD1L* could be also implicated in the general height abnormalities observed in 1q21.1 distal CNV carriers. An effect on height has been reported for the distal deletion, with 25–50% of the carriers having short stature, whereas the duplication carriers tend to be in the upper percentiles of height(4, 13, 14). *CHD1L* overexpression led to a significant increase of the body length (**Figure 1J and K**) and inter-somites distance (**Supplementary Figure S1C and D**) at 5 dpf in zebrafish larvae, supporting the possibility that *CHD1L* could broadly impact anthropometric traits and growth.

To validate further the specificity of the phenotypes driven by *CHD1L* and to investigate whether it was possible to simulate the reciprocal phenotypes seen in individual carriers of 1q21.1 distal deletion, we sought to determine the effect of a loss of the sole ortholog of *CHD1L* in zebrafish. *chd1l* is ubiquitously expressed as early as 3 hours post-fertilization (hpf) in blastomeres with an expression peak between 3 and 72 hpf (Daniocell database; **Supplementary Figure S2A and B**). More specifically, *chd1l* is expressed in all brain vesicles including telencephalon, diencephalon, midbrain, midbrain-hindbrain boundary and neural crest cells between 12 and 48 hpf. These expression data confirmed the relevance of a zebrafish model to study the developmental role of CHD1L. We thus obtained an *N*-ethyl-*N*- nitrosourea (ENU) mutagenized stable line for the sole *chd1l* zebrafish orthologue (*chd1l^sa14029^* line carrying a p.Arg6X stop mutation) and we sought to determine the total brain volume of 5 dpf *chd1l^+/+^*and *chd1l^-/-^* larvae using High-Resolution Episcopic Microscopy(40). We detected a ∼25% reduction of the brain in *chd1l^-/-^* larvae compared to wild-type larvae (**Figure 2A and B**). Bi-dimensionally measurements confirmed a significant decrease of head size (**Figure 2C and Supplementary Figure S1E**) and body length (**Figure 2D and Supplementary Figure S1F**) in homozygous *chd1l* mutant larvae at 5 dpf. Importantly, these phenotypes are specific to *chd1l*; both morphometric phenotypes could be fully rescued upon injection of 100 pg of wildtype human *CHD1L* mRNA in the *chd1l^-/-^* homozygous eggs at 1-2 cell stage (**Figure 2C and D and Supplementary Figure S1E and F**). There was also no defect in other structures, including the heart and the swim bladder, indicating the absence of global developmental delay of the *chd1l* mutant larvae. Notably, the reduction of the head size and body length measurements at 5 dpf was also observed in *chd1l*^+/-^ heterozygous mutant larvae, which mimicked the gene dosage of heterozygous human 1q21.1 distal deletion (**Supplementary Figure S3A-D**).

**Figure 2:**
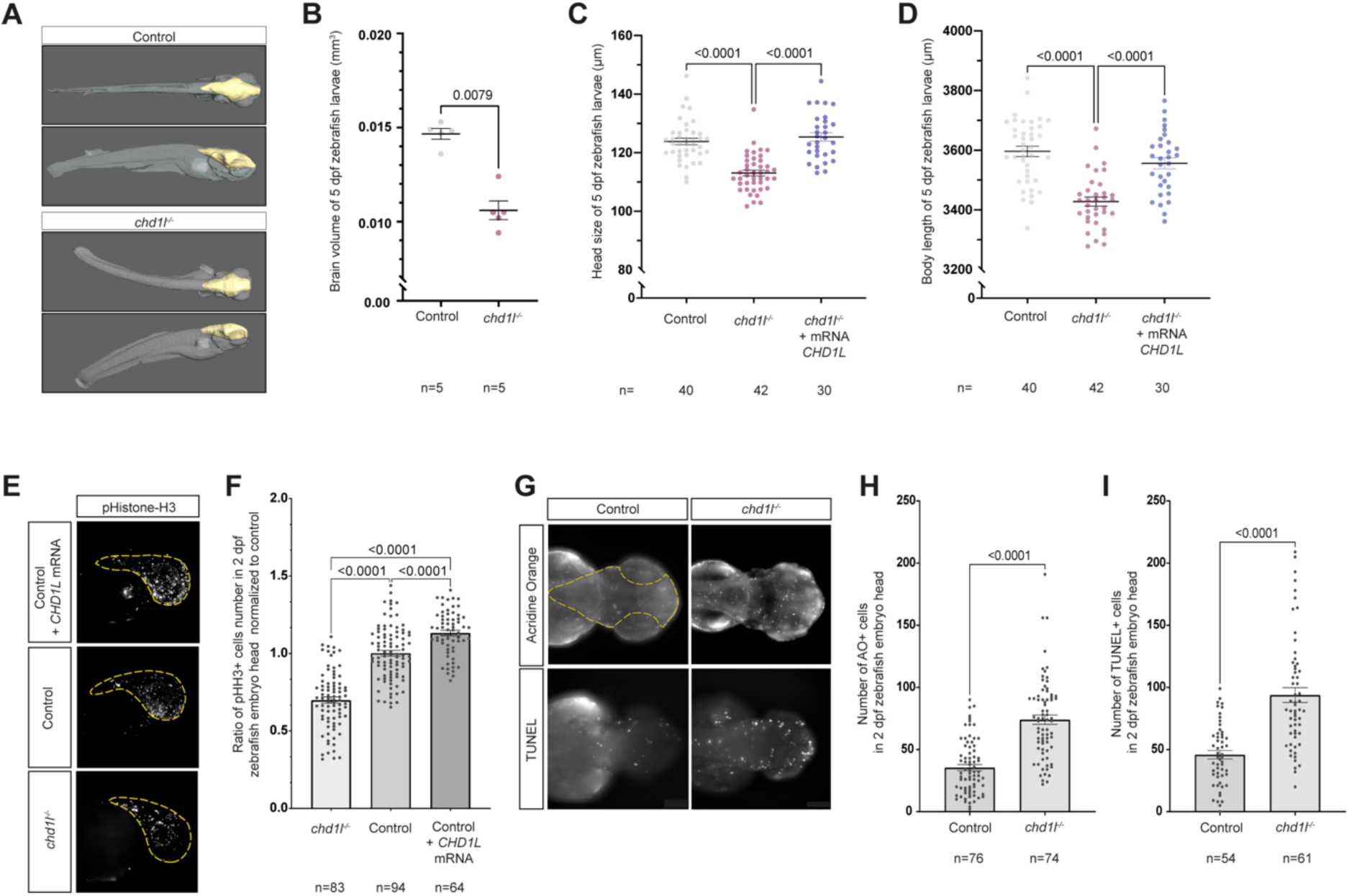
Loss of *chd1l* recapitulates mirrored neuroanatomical phenotypes of the 1q21.1 distal deletion in zebrafish. **(A)** 3-Dimensional reconstitution of 5 dpf control and *chd1l*^-/-^ zebrafish larvae acquired by HREM. The brain volume is shown in yellow. **(B)** Dot plot showing the quantification of brain volume of 5 dpf control and *chd1l^-/-^* zebrafish larvae. Data shown as mean ± SEM of a representative batch; Student’s *t* test. **(C-D)** Dot plot showing the rescue of *chd1l^-/-^* phenotypes for head size and body length upon injection of *CHD1L* mRNA (100 pg) in *chd1l ^-/-^* larvae compared to control larvae at 5dpf respectively. Data shown as mean ± SEM of triplicate batches; Ordinary One-way ANOVA. **(E)** Lateral views of 2 dpf zebrafish heads for mutant *chd1l^-/-^*, control and *CHD1L* mRNA-injected larvae stained with phospho-histone H3 (pHH3; M-phase marker). Dotted yellow lines indicate quantified head areas. **(F)** Dot plot showing the pHH3 ratio of proliferating cells in the heads of 2 dpf *chd1l^-/-^* and *CHD1L* mRNA-injected larvae normalized to control. Data shown as mean ± SEM of triplicate batches; Ordinary One-way ANOVA. **(G)** Dorsal views of 2 dpf heads of control and *chd1l^-/-^* mutant larvae stained with Acridine Orange (AO) or TUNEL. Dotted yellow line indicates quantified head area. **(H)** Dot plot showing the number of AO-positive cells in the heads of 2 dpf control and *chd1l^-/-^* mutant larvae. Data shown as mean ± SEM of triplicate batches; Mann-Whitney test. **(I)** Dot plot showing the number of TUNEL-positive cells in the heads of 2 dpf control and *chd1l*^-/-^ mutant larvae. Data shown as mean ± SEM of triplicate batches; Mann-Whitney test. To ease the visualization of the data, the non-significant *p*-values are not showed on the graphs.

### *CHD1L* dosage perturbation affects cell proliferation and cell death

To investigate the mechanism leading to the head size differences, we examined the developing brain in both *CHD1L* overexpressant and *chd1l^-/-^* mutant embryos. First, we determined the number of mitotic cells by phospho-histone H3 immunostaining of *chd1l^-/-^*, control and *CHD1L* mRNA-injected zebrafish heads at 2 dpf (**Figure 2E**). A significant decreased number of proliferating cells in the head at 2 dpf was observed in *chd1l^-/-^* mutant larvae compared to controls. Conversely, an increased cell proliferation in the head at 2 dpf was observed upon *CHD1L* overexpression (**Figure 2F**). Second, we assessed cell viability by performing Acridine Orange staining and TUNEL assay on control and *chd1l*^-/-^ larvae at 2 dpf (**Figure 2G**). Both staining revealed that loss of *chd1l* increases apoptotic level in the head of the larvae, supporting the microcephalic phenotype observed at 5 dpf (**Figure 2H and I**). Such imbalance of cell proliferation and apoptosis have been previously seen in zebrafish and mice modeling the micro/macrocephaly phenotypes(68–71).

Although the zebrafish brain bears several similarities with the mammalian brain in terms of developmental programming, we examined whether our findings might be relevant to cortical development in a mammalian system. In both human and mouse, *CHD1L/Chd1l* is ubiquitously expressed; we noted similar profiles in both species in the whole brain with an expression peak at 4 weeks post-conception in human and murine E11.5 (**Supplementary Figure S2C and D**)(72). We performed *in utero* electroporation (IUE) to overexpress or deplete *Chd1l* in mouse embryonic cortices and investigated the potential of apical progenitors lining the lateral ventricle to generate TBR1- positive cortical neurons. We performed IUE of plasmids expressing mouse *Chd1l* transcripts or siRNA directed against mouse *Chd1l* together with a pCAGGS-GFP reporter construct, allowing the expression of GFP specifically in electroporated apical progenitors and their progeny, in wild-type mouse cortices at E13.5 (**Figure 3A**). One day after electroporation, we evaluated the number of TBR1-positive neurons produced from the electroporated progenitors. When compared to respective controls (empty vector, or siRNA directed against luciferase), overexpression of *Chd1l* induced a significant increase (+19.7%) of TBR1-positive neuronal progeny (**Figure 3B and C**) whereas depletion of *Chd1l* impaired the generation of TBR1-positive cortical neurons (-24%) (**Figure 3D and E**). These mirrored phenotypes in mouse cortices strongly showed that function of *Chd1l* during neuronal development is conserved between non-mammalian vertebrates and mammals and that the dosage of *Chd1l* is critical for neuronal production.

**Figure 3:**
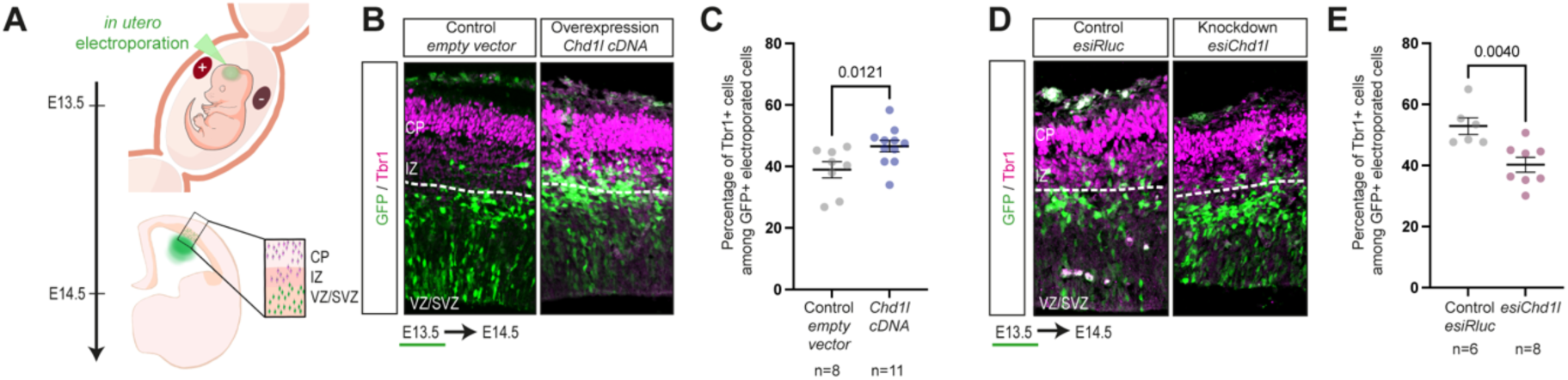
Dosage changes of *Chd1l* controls neuronal production in transient mouse model. **(A)** Schematic of *in utero* electroporation. Mouse embryos are electroporated at E13.5, mouse brain are dissected at E14.5 and stained for Tbr1 neuronal marker. Double GFP-positive and Tbr1-positive cells representing the newborn neurons are counted in a defined area in the VZ/SVZ. **(B)** E14.5 mouse brain slices for control and *Chd1l* overexpressant conditions. **(C)** Dot plot showing the percentage of Tbr1-positive cells among the electroporated GFP-positive cells from the defined area in E14.5 mouse brains from control or electroporated with *Chd1l* expression vector. Data shown as mean ± SEM, electroporated and imaged embryos were from five different litters for control condition and from six different litters for *Chd1l* overexpression condition; Student’s t-test. **(D)** E14.5 mouse brain slices for control and knockdown *Chd1l* conditions. **(E)** Dot plot showing the percentage of Tbr1-positive cells among the electroporated GFP-positive cells in E14.5 mouse brains from control or electroporated with esi*Chd1l* RNA. Data shown as mean ± SEM, electroporated and imaged embryos were from three different litters for *esiRluc* condition and from four different litters for *esiChd1l* condition; Student’s *t*-test. CP, cortical plate ; IZ, intermediate zone ; VZ/SVZ, ventricular/subventricular zones.

Taken together, the combination of our zebrafish and mouse data revealed that *CHD1L* is a major contributor of the mirrored body size and head size phenotypes associated with the 1q21.1 distal CNV. However, our data do not exclude the possibility that other loci within the region also have an independent contribution to the 1q21.1 distal deletion or duplication anatomical phenotypes. Therefore, we performed the combinatorial overexpression of *CHD1L* with each of the remaining 1q21.1 distal transcripts to evaluate whether other genes within the CNV might also be relevant to the head size phenotypes through additive or multiplicative interactions with *CHD1L*. None of the other transcripts exacerbated nor alleviated the macrocephaly phenotype driven by *CHD1L* (**Supplementary Figure S4A**), indicating that none of the other genes within this interval act as modifier for the head size phenotype, at least as determined by our assays.

### Loss of *CHD1L* disrupts neurogenic program in human neuronal progenitor cells

To our knowledge, *CHD1L* has never been shown to control human neurogenesis. Therefore, to investigate *CHD1L*’s potential gene targets and associated pathways in the neuro-developmental context, we designed a guide RNA targeting exon 1 and performed CRISPR-Cas9 editing of *CHD1L* in human induced pluripotent stem cells (hiPSC GM8330-8, used in(73)) adapting the protocol described in(74). We selected two isogenic hiPSCs lines carrying the following homozygous frameshift mutations, c.72_76delCCGAG (hiPSC *CHD1L*^-/-^ line 1) and c.75_76delAG (hiPSC *CHD1L*^-/-^ line 2) (**Supplementary Figure S5A**), respectively, and confirmed the absence of further mutations in the Top-20 predicted off-targets (**Supplementary Table S1**). Karyotype analysis confirmed the genomic integrity of CRISPR-generated lines as well as control hiPSCs and allowed to exclude the presence of aneuploidies and large chromosomal rearrangements that could have been introduced during the gene-editing process (**Supplementary Figure S6A-C**). In parallel, we confirmed the absence of the *CHD1L* protein in both lines by western blot (**Figure 4A**). Given that recent studies on brains from ASD individuals highlight transcriptional changes occurring primarily in neuronal lineages, we chose to focus our analyses on neuronal progenitor cells, where these changes are first observed (75, 76). The control and the *CHD1L*- edited hiPSC lines were thus differentiated into human neural progenitor cells (hNPC) (**Figure 4B**). After seven days of differentiation, formation of rosettes was observed. Positive expression of hNPC markers SOX1 and PAX6 confirmed the lineage commitment (**Figure 4C**) and validated the successful differentiation into hNPC. RNA-seq analysis of the derived cells confirmed that both mutant lines shared similar expression profiles (**Figure 4D**). Further, we intersected the list of differentially expressed genes (DEG) from both mutant lines and we found a total of 857 common DEG in hNPC lacking *CHD1L* compared to control hNPC, including 563 down-regulated and 294 up-regulated genes (FDR <0.05; **Figure 4E and Supplementary Table S2**).

**Figure 4:**
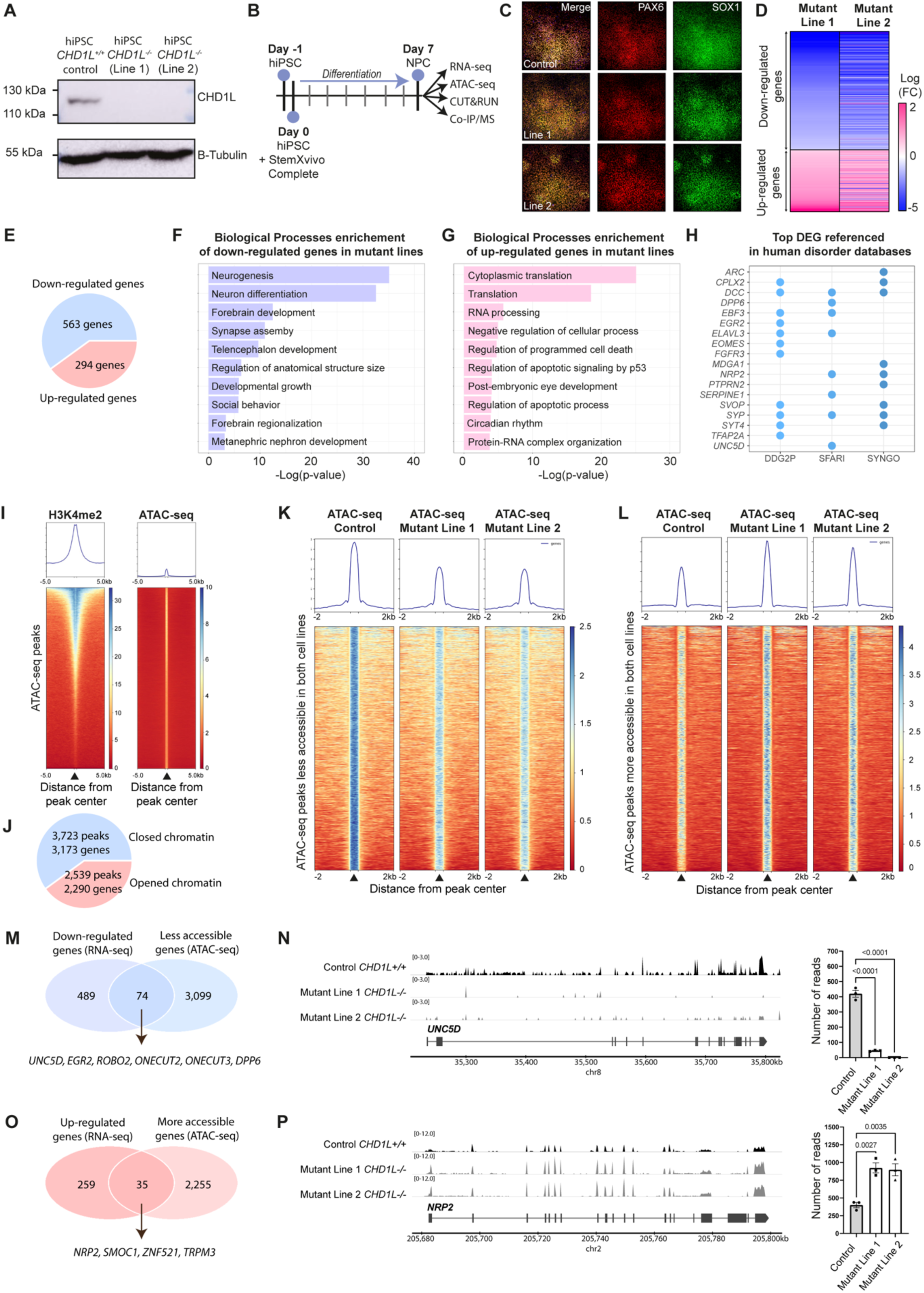
Loss of *CHD1L* perturbs expression and chromatin accessibility of neurogenesis-associated genes in hNPC. **(A)** Western blot analysis of CHD1L protein level in control hiPSC *CHD1L^+/+^* and in the two isogenic *CHD1L^-/-^* cell lines generated by CRISPR-Cas9 editing showing the absence of CHD1L protein in mutant lines. **(B)** Schematic representation of human Neural Progenitor cells (hNPC) generation from hiPSC and subsequent analysis pipeline. **(C)** Immunostainings of hiPSC-derived hNPC with PAX6 and SOX1 neural progenitor markers. **(D)** Heatmap of the differentially expressed genes in hNPC Mutant Line 1 compared to Mutant Line 2 showing similar expression profiles between both mutants. **(E)** Pie chart of differentially expressed genes in *CHD1L^-/-^* hNPC compared to control hNPC (FDR<0.05). **(F and G)** Gene Ontology Biological Processes enrichment of the downregulated and upregulated genes in mutant hNPC respectively. **(H)** Dot plot of referenced genes in indicated databases associated with human disorders and synaptogenesis function among the 52 most DEG (|Log2FC|> 1; FDR <0.05) in *CHD1L* mutant hNPC. **(I)** Average tag density profiles and corresponding heatmap representation of tag density map of H3K4me2 CUT&RUN peaks, ± 2 kb from peak center of ATAC-seq peaks from control hNPC. **(J)** Pie chart of differentially accessible chromatin sites in *CHD1L^-/-^* mutant hNPC (Line 1 and Line 2 combined) compared to control hNPC. **(K)** Tag density map of control, mutant Line 1 and mutant Line 2; ± 2 kb from peak center for 3,723 less accessible and 2,539 more accessible peaks in *CHD1L*^-/-^ hNPC and corresponding average tag density profiles. Line 1 and Line 2 were combined and the same direction of change for both mutants was selected; Mixed refer to peaks that exhibit opposite changes in mutant Line 1 compared to mutant Line 2 and were discarded from the subsequent analyses. **(L)** Venn diagram showing 74 down-regulated genes that are also less accessible in absence of *CHD1L*. **(M)** Chromatin accessibility and level of expression for *UNC5D* in control and mutant hNPC. **(N)** Venn diagram showing 35 up-regulated genes that are also more accessible in absence of *CHD1L*. **(O)** Chromatin accessibility and level of expression for *NRP2* in control and *CHD1L* mutant hNPC.

To interpret the transcriptomic changes, we conducted a pathway analysis on the list of the 563 downregulated genes using a PANTHER overrepresentation test (geneontology.org) for biological processes. The analysis revealed significant enrichment of terms associated to neurodevelopment and maturation processes in down-regulated genes: “Neurogenesis” (GO:0022008, FDR = 3.16E-32), “Forebrain development” (GO:00309005, FDR = 1.02e-10); “Forebrain regionalization (GO: GO:0021871, FDR = 1.94e-2)”; “Synapse assembly” (GO:0007416, FDR = 2.94e-9); “Developmental growth” (GO:0048589, FDR = 1.34e-4) and terms associated to renal development “Metanephric nephron development” (GO: GO:00722721, FDR = 3.07e-2) (**Figure 4F**). Conversely, up-regulated genes are associated to various translational processes such as “Translation” (GO:0006412, FDR = 3.04e-19) and “RNA processing” (GO:0006396, FDR = 1.83e-6) in addition to cell death processes “Regulation of programmed cell death” (GO:0043067, FDR=1.8e-5), “Regulation of apoptotic signaling by p53” (GO:1902253, FDR=8.64e-5). We also noted association to the “Post-embryonic camera-type eye development” (GO:0031077, FDR = 3.09e-2) as the most enriched term (**Figure 4G and Supplementary Table S2**). Further, analysis of the reactome pathways confirmed the association of down-regulated genes to the nervous system and nephrotic development whereas up-regulated genes tend to be associated to DNA damage response and cell cycle in absence of *CHD1L* (**Supplementary Figure 5B and C and Supplementary Table S2**).

To determine the gene sets that are the most affected by *CHD1L* loss, we applied an absolute value of |Log2FC|> 1 and we obtained 52 DEG including 45 down-regulated and 7 up-regulated genes between control and both mutant hNPC lines (**Supplementary Table S2**). We constructed a functional protein-protein interaction network of the identified 52 DEG using STRING(60) (string-db.org), which highlighted the presence of a significantly greater number of protein-protein interactions than expected (*p*=1.0*×*10^−16^) with three functional clusters associated to brain regionalization/nervous system development, regulation of neurotransmitter levels and axonogenesis-associated terms (**Supplementary Figure S5D**). Furthermore, 8/52 genes (*DCC*, *DPP6, EBF3, ELAVL3, NRP2, SERPINE1, SYP, UNC5D*) were found in the Simons Foundation Autism Research Initiative (SFARI) gene list for their association to autism spectrum disorders (enrichment *p*=0.02), and a significant enrichment was also identified for the developmental disorder genes (11/52 genes: *CPLX2*, *DCC*, *EBF3*, *EGR2*, *ELAVL3*, *EOMES*, *FGFR3*, *SVOP*, *SYP*, *SYT4*, *TFAP2A*) in the Decipher Database (*p*=6e-4). A total of 9/52 genes are referenced in the SynGO database recapitulating genes associated to synapse (*ARC*, *CPLX2*, *DCC*, *MDGA1*, *NRP2*, *PRPTRN2*, *SVOP*, *SYP* and *SYT4*) (*p*=0.001) (**Figure 4H**). One more autism risk DE gene (*THSD7A*) was annotated by crossing our list with the *de novo* “high confidence” ASD genes carrying likely gene disrupting mutations(77); interestingly, two pleiotropic genes (*DCC, UNC5D*) with a prominent role in different psychiatric disorders (78) were also identified among the 52 DEG. A trend for enrichment was found when we intersected the 52 DEG with a list of manually curated growth genes obtained from two publications(79, 80) (4 genes: *DPP6, FGFR3, TFAP2A, IGFBP3*). Remarkably, the long noncoding RNA *PAX8-AS1* associated with hypothyroidism(81) was one of the most down- regulated in absence of *CHD1L* (Log2FC <-4). Moreover, other genes associated with hypothyroidism and short stature were found downregulated (e.g. *SOX3, GLI2, KMT2D*, and *FGD1*)(82–88) (**Supplementary Table S2**). Of note, individuals with 1q21.1 deletion often exhibit short stature(13, 14) and one case has been reported with congenital hypothyroidism(89).

### Chromatin accessibility of neurodevelopmental-associated genes is altered in *CHD1L*-depleted human neuronal progenitor cells

CHD1L is a known ATP-dependent chromatin remodeler(28–34). To determine whether the transcriptomic changes are due to perturbed chromatin accessibility, we performed Assay for Transposase-Accessible Chromatin with high throughput sequencing (ATAC-seq) on control and *CHD1L^-/-^* mutant hNPC lines (48). We validated that the detected ATAC-seq peaks from control cell line colocalized with the presence of dimethylated lysine residue at position 4 of histone H3 (H3K4me2), a hallmark for active promoters and enhancers **Figure 4I and Supplementary Figure S5A and B**). In the absence of *CHD1L*, we found a total of 3,723 peaks for which the chromatin was less accessible and 2,539 peaks that were more accessible in hNPC (**Figure 4J and K**), further demonstrating that *CHD1L* plays a role in chromatin remodeling in neuronal progenitors (**Figure 4K and Supplementary Table S3**). We next intersected the DEG with the list of differentially accessible genes detected in *CHD1L^-/-^* mutant hNPC. We found a total of 74 genes that were coordinately less expressed and showed closed chromatin sites at their loci as exemplified for ASD and ADHD-susceptibility gene *UNC5D*(90–94) (**Figure 4L and M**), and 35 genes that were coordinately more expressed and were associated to newly accessible chromatin sites as seen for *NRP2* that regulates dendritic growth of adult-born neurons in the dental gyrus(95, 96) (**Figure 4N and O and Supplementary Table S3**).

Next, TOBIAS ATAC-seq footprinting analysis(50) unraveled that 19 clusters of transcription factors had favorized binding abilities when *CHD1L* was lost, whereas 8 clusters were more able to bind chromatin when *CHD1L* was expressed ((FDR<0.01, **Figure 8A and B**). The 27 clusters included 551 individual transcription factors (**Supplementary Table S3**). In absence of *CHD1L*, we noted that the binding motifs of POU5F1, SOX2, CTCF, FOXO1, FOXP2 were more accessible whereas the binding motifs for GLI2, GLI3, JUN, SMAD2::SMAD3::SMAD4, TP53 were less accessible (**Figure 8C and Supplementary Table S3**). Most of those factors are found to be dysregulated in ASD brain tissues(75).

Taken together, the transcriptomic data and footprinting analysis collectively revealed that the transcriptional perturbation in the *CHD1L*-edited hNPC lines specifically impairs cell identity, brain regionalization, neurogenesis, synaptogenesis and growth-associated pathways. Notably, the most affected genes are linked to ASD and developmental disorders. These disrupted pathways align with known signatures of ASD identified through both bulk and single-cell genomic studies on tissues from ASD individuals, reinforcing the notions that CHD1L plays a critical role in neurodevelopment and that its dosage imbalance likely underlies the cognitive impairments associated with 1q21.1 CNV.

### CHD1L acts as a co-transcription factor interacting with NuRD complex and the pioneer transcription factor SOX2

To identify direct CHD1L targets, hNPC chromatin was immunoprecipitated with antibody against CHD1L, followed by massive parallel sequencing using CUT&RUN(97). Bioinformatics analysis uncovered 5,070 CHD1L peaks, mainly located at intronic and intergenic regions (**Figure 5A and Supplementary Table S4**). These peaks were assigned to 4,002 unique genes (**Figure 5B and Supplementary Table S4**). Amongst these genes with CHD1L peaks, we found 102 genes that were down-regulated including key neurodevelopmental genes such as *ONECUT2* and *ONECUT3* and 55 genes that were up-regulated in our transcriptome data (**Figure 5C and Supplementary Table S4**). An analysis of genomic distribution revealed that CHD1L peaks did not correlate with H3K4me2 peaks and were present in far upstream and downstream regions (−100 to −50 kb and 50 to 100 kb), intermediate locations (−50 to −20 kb and 20 to 50 kb) and DNA segments in the vicinity of the transcription start site (TSS) (−10 to 10 kb) (**Figure 5D and E**).

**Figure 5:**
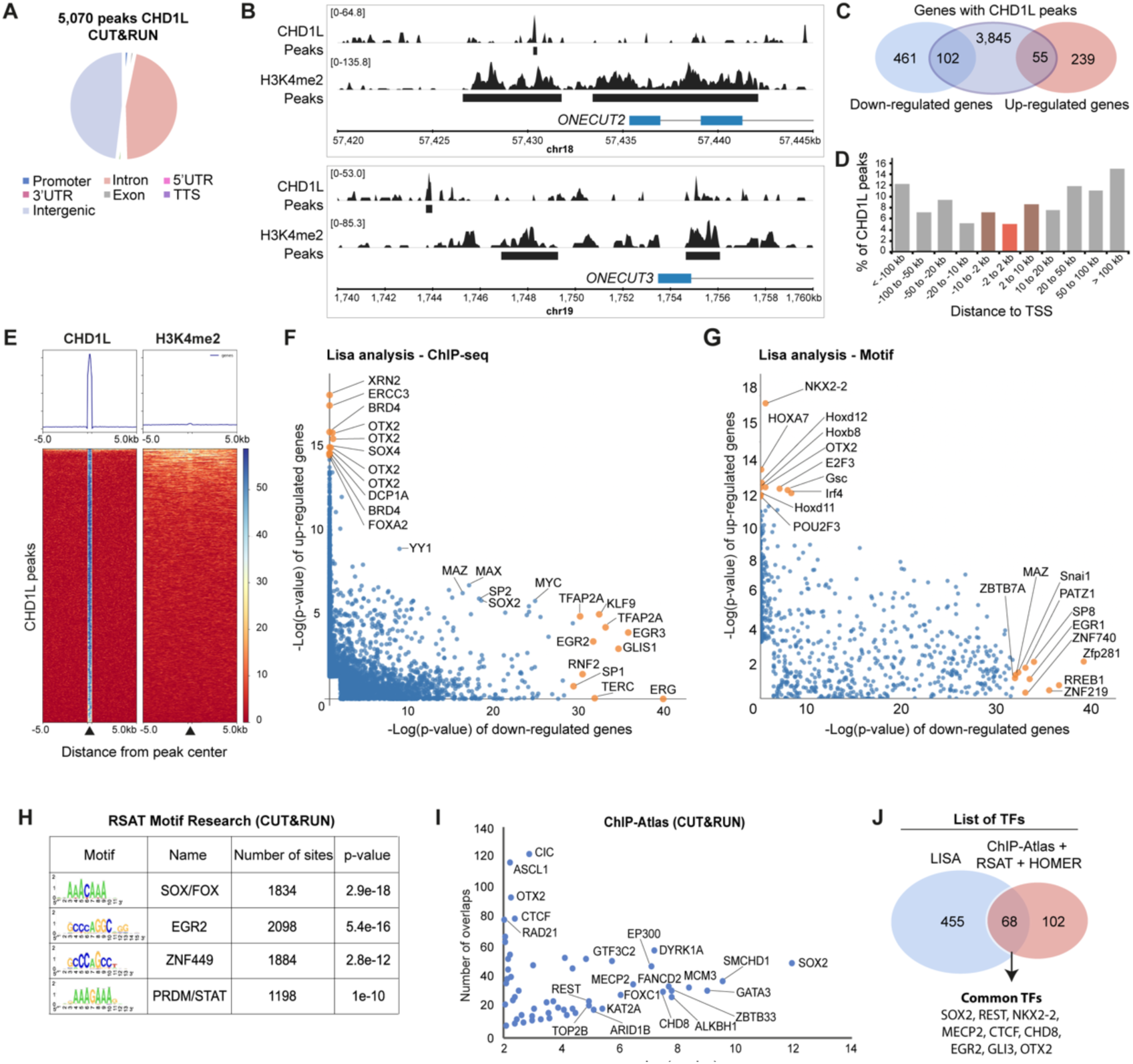
Characterization of CHD1L binding in hNPC. **(A)** Pie charts of the genomic distribution of CHD1L CUT&RUN peaks in control hNPC. **(B)** Examples of CHD1L-bound sites located upstream of down-regulated genes *ONECUT2* and *ONECUT3*. **(C)** Venn diagram showing CHD1L-bound genes that are down- or up-regulated in *CHD1L* mutant hNPC. **(D)** CHD1L Peak distribution from TSS. **(E)** Tag density map of CHD1L and H3K4me2 CUT&RUN peaks, ± 2 kb from the CHD1L peak center and corresponding average tag density profiles. **(F and G)** Lisa results showed as pairwise scatterplots for comparison of the two gene sets (down- and up-regulated genes in absence of *CHD1L* in hNPC) based on transcription factor ChIP-seq and transcription factor motifs respectively. The top 10 transcription factors are shown in yellow. **(H and I)** RSAT-based motif enrichment analysis and ChIP-Atlas comparison of *CHD1L* CUT&RUN data respectively. **(J)** Venn diagram showing common transcription factors predicted by Lisa, ChIP-Atlas, RSAT and HOMER analyses.

To identify putative transcription factors (TFs) that may be involved in the recruitment of CHD1L during neurogenesis, we predicted the transcriptional regulators of the DEG in *CHD1L^-/-^*hNPC by inferring transcriptional regulators through integrative modeling of public chromatin accessibility and ChIP-seq data (http://lisa.cistrome.org/)(98). Notably, the Top-10 predicted regulators of the up-regulated genes in *CHD1L^-/-^* NPC included transcription factors involved in DNA damage response and repair (*XRN2, ERCC3, SOX4, BRD4*) and in early morphogenesis of the central nervous system (*OTX2*). We also noted a strong enrichment of *NKX2.2* binding motif, that is known to control ventral neuronal patterning(99) and to regulate oligodendrocyte differentiation(100). For the down-regulated gene set, cistromic analysis highlighted the EGR family of transcription regulatory factors (*EGR1, EGR2*), which is implicated in orchestrating the changes in gene expression that underlie neuronal patterning and plasticity(101, 102), the architectural protein CTCF that changes the higher-order chromatin structure and controls the distance between associating domains within and among chromosomes, and several TFs controlling pluripotency and differentiation such as SOX2, GLIS1, ZFP281, and ERG(103–106) (**Figure 5F and G and Supplementary Table S2**). We next performed RSAT and HOMER motif analyses as well as ChIP-Atlas comparisons on CHD1L CUT&RUN data, and found a strong enrichment for SOX/FOX, EGR2, ZNF449 and PRDM/STAT binding sequences in hNPC (**Figure 5H and I and Supplementary Table S4**). By intersecting the lists of transcription factors from the different prediction tools utilized on our transcriptomic and cistromic data, we identified 68 candidate transcription factors including SOX2, OTX2, MECP2, CTCF, CHD8, EGR2 and GLI3 (**Figure 5J and Supplementary Table S4**). This list suggested the implication of CHD1L in major neuro-developmental pathways controlling proper nervous system development (SOX2, OTX2, EGR2, GLI3, CTCF)(102, 107–112) and with genes mutated in neurodevelopmental disorders (CHD8, MECP2)(69, 113).

To identify protein partners of CHD1L in hNPC and whether the aforementioned predicted TFs interact physically with CHD1L, we performed affinity purification with CHD1L antibody followed by mass spectrometry (AP-MS) on both wildtype and *CHD1L*^-/-^ mutant lines. By applying stringent thresholds, we identified a total of 286 candidate partners (**Supplementary Table S5**). To test whether some TFs were present in the AP-MS dataset, we filtered the candidates based on their cellular localization, and we retrieved 104 nuclear partners of CHD1L. STRING analysis revealed four functional clusters including DNA-repair proteins (Cluster 1), components of the NuRD complex (Cluster 2), Arp2/3 complex proteins (Cluster 3) and Histone domains (Cluster 4). More specifically, Cluster 2 included core components of the NuRD complex (GATAD2, MBD3, HDAC1, HDAC2, MTA2 and RBBP4), the ATP-dependent remodeling factor CHD7, and the transcription factors SOX2, OTX2 and PAX6 known for their role in cell-fate and brain regionalization(107, 114, 115) (**Figure 6A and B and Supplementary Table S5**). Cluster 2 and 4 also includes candidate protein partners associated to cell cycle and cell survival processes (“G1/S specific transition”, “Apoptosis induced DNA fragmentation”) (**Figure 6B and Supplementary Table S5**).

**Figure 6:**
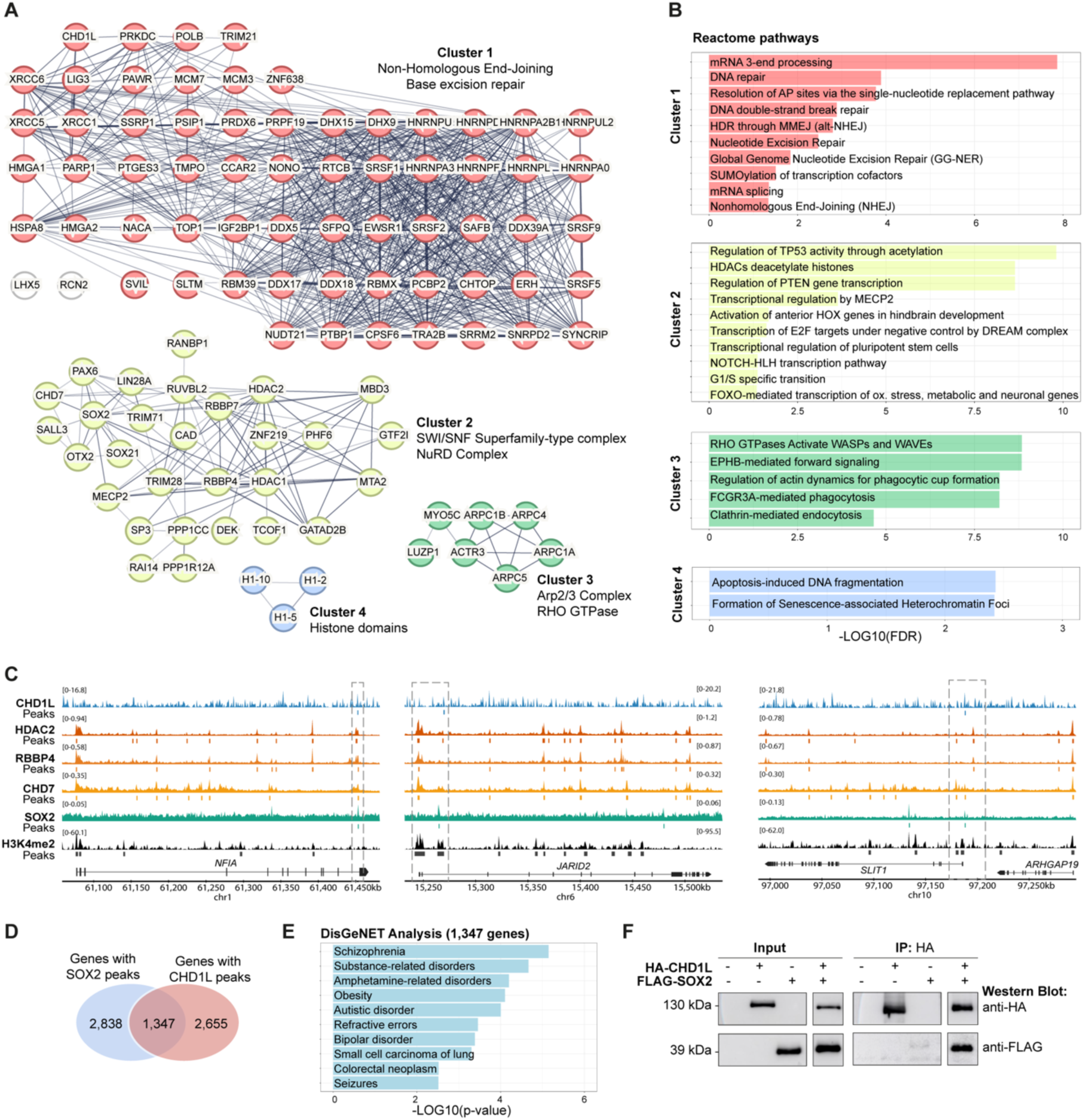
Immunoprecipitation and mass-spectrometry define CHD1L interactome and reveal developmental partners in hNPC. **(A)** STRING analysis of 104 nuclear proteins co-immunoprecipitated with CHD1L in hNPC extracts. Protein-Protein interactions are shown (PPI enrichment *p*-value <1e^-16^) and annotated according to functional clusters: Cluster 1 (63 proteins), Cluster 2 (29 proteins), Cluster 3 (7 proteins), Cluster 4 (3 proteins). **(B)** Gene ontology analyses showing reactome pathways enriched for each cluster. **(C)** Representative co-localization of CHD1L, HDAC2 (SRX19212560), RBBP4 (SRX19212562), SOX2 (SRX330107), CHD7 (SRX9795022) and H3K4me2 peaks on neuro-developmental genes (*NFIA, JARID2, SLIT1*) in human neural cells. **(D)** Venn diagram of SOX2 target genes (SRX330107) and CHD1L target genes showing an overlap of 1,347 genes. **(E)** DisGeNET analysis of the common 1,347 SOX2 and CHD1L target genes showing enrichment in human neurodevelopmental disorders. **(F)** Representative western blot of CHD1L and SOX2 co-immunoprecipitation in transfected HEK cells.

Taking advantage of available ChIP-seq datasets available, we observed co-localization of HDAC2, RBBP4 (NuRD complex components), CHD7, SOX2 and CHD1L peaks on neurodevelopment-associated genes such as *NFIA*, *JARID2* and *SLIT1* further suggesting that CHD1L could act with the NuRD complex to regulate gene expression (**Figure 6C**). Using SOX2 ChIP-seq compared to our CHD1L CUT&RUN, we found that one third of genes bound by CHD1L were also bound by SOX2 in human neuronal progenitor cells (**Figure 6D and Supplementary Table S4**). Remarkably, DisGeNET analysis on the 1,347 common genes revealed an enrichment for terms including schizophrenia, autistic disorder, obesity and seizures (**Figure 6E**). Co-immunoprecipitation experiments further confirmed the physical interaction between SOX2 and CHD1L (**Figure 6F**). Overall, our data strongly indicate that CHD1L could affect gene expression through its direct or indirect interaction with pioneer TFs and NuRD regulatory complex during human neurodevelopment and brain regionalization.

### CHD1L loss perturbs cell-fate decision during forebrain regionalization

Self-organizing cerebral organoids grown from pluripotent stem cells combined with single-cell genomic technologies provide opportunities to examine gene regulatory networks underlying human brain development and diseases (116–118). To better characterize the role of *CHD1L* during cerebral development and to test the possibility that *CHD1L* plays a role during brain regionalization as indicated by our omics data, we derived *CHD1L^+/+^* and *CHD1L ^-/-^* hiPSC (mutant Line 1) into cerebral organoids for 60 days *in vitro* (DIV). We chose to build organoids through a self-organization process by providing a permissive environment with minimal external cues which allowed us to determine the intrinsic capacity of the control and *CHD1L* mutant hiPSC to undergo *in vivo*-like morphogenesis(117) (**Figure 7A**). *CHD1L^-/-^* (Line 1) exhibited apparent similar morphology as *CHD1L^+/+^*control organoids during the *in vitro* maturation (**Figure 7B**). At 52 DIV, control and mutant organoids were positive for both SOX2 (neural progenitor cells marker) and TUJ1 (neural cytoskeleton marker) (**Figure 7C**). We noted typical structures of cerebral organoids including rosettes (SOX2+) and cortical plate layers (TUJ1+)(118). However, we observed a decreased level of TUJ1 protein in *CHD1L*^-/-^ mutant organoids compared to control organoids at 60 DIV (**Figure 7D and Supplementary Figure S9A**), suggesting impaired neurogenesis in organoids lacking *CHD1L*.

**Figure 7:**
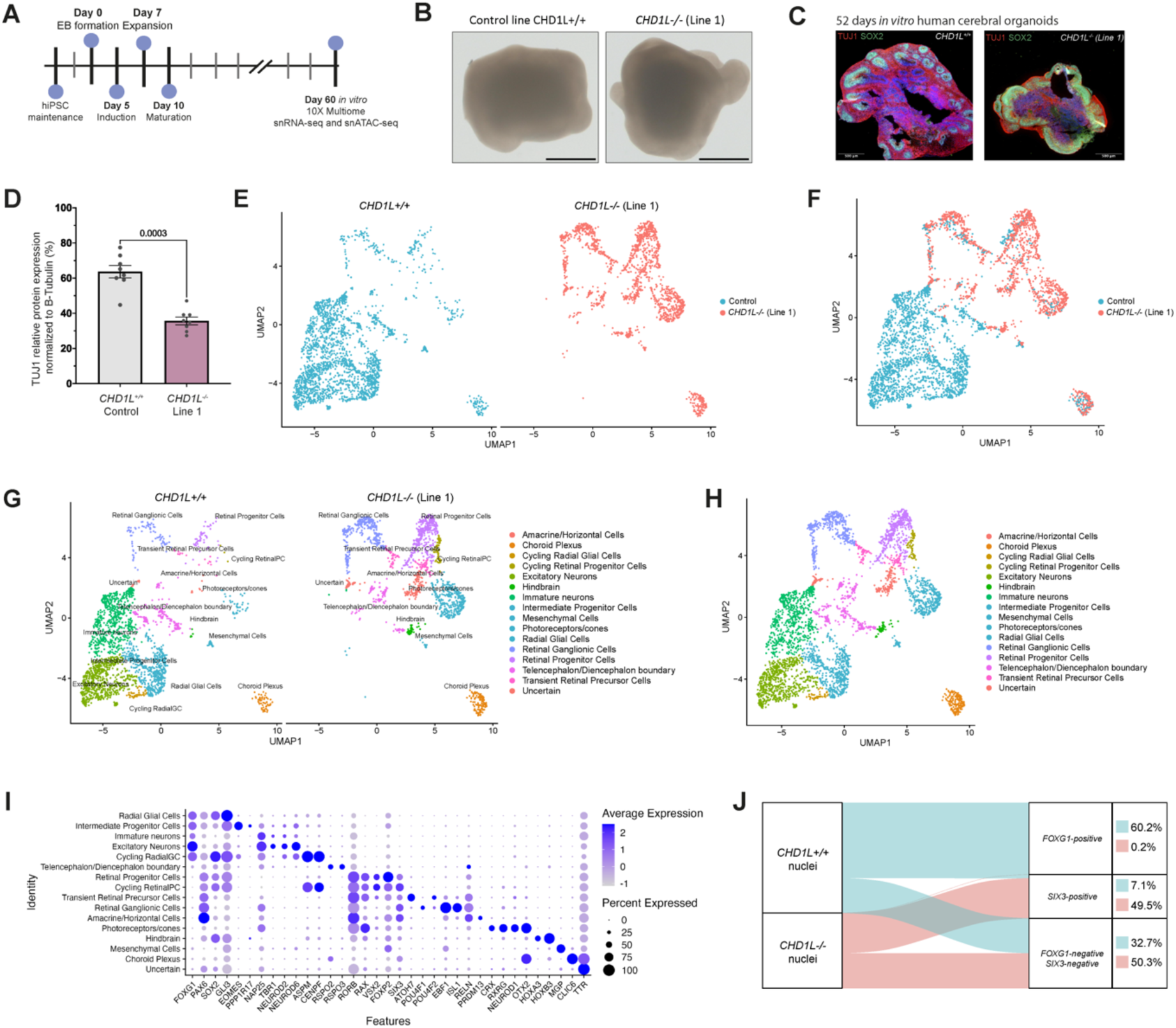
Loss of CHD1L perturbs forebrain cell-fate determination in human cerebral organoids. **(A)** Schematic of human cerebral organoid (hCO) generation from hiPSC. **(B)** Pictures of 60 days *in vitro* (DIV) hCO derived from *CHD1L*^+/+^ and the two *CHD1L*^-/-^ cell lines. Scale bar, 1 mm. **(C)** Immunostaining against TUJ1 (neuronal cytoskeleton marker) and SOX2 (neural progenitor cells) on 52 DIV hCO either *CHD1L^-/-^* (mutant Line 1) or control. **(D)** Semi-quantitative analysis of TUJ1 protein expression in *CHD1L^-/-^* (mutant Line 1) and control hCO normalized with β-Tubulin. Data shown as mean ± SEM of 8 hCO per condition; representative of N=2 batches of generated organoids; Student’s *t*-test. **(E)** UMAP projection of *CHD1L*^+/+^ and *CHD1L*^-/-^ hCO cells colored by condition. N=6 organoids per condition, 2,385 cells *CHD1L*^+/+^ and 1,634 cells *CHD1L*^-/-^ sequenced. **(F)** UMAP representation of *CHD1L*^+/+^ and *CHD1L*^-/-^ (mutant Line 1) multiome analysis combined and integrated on the same graph. N=12 organoids, 4,019 cells. **(G)** UMAP representation of *CHD1L*^+/+^ and *CHD1L*^-/-^ (mutant Line 1) hCO multiome analysis on the same graph. Clusters based on cell identities are indicated. **(H)** UMAP representation of *CHD1L*^+/+^ and *CHD1L*^-/-^ (mutant Line 1) hCO multiome analysis combined and integrated on the same graph. Clusters based on cell identities are indicated. **(I)** Dot plot representing key cell type markers and their levels of gene expression within cerebral organoids’ cell clusters. Dot size indicates the proportion of cells per cluster expressing the corresponding gene and color is associated to the average expression level of the corresponding gene per cluster. **(J)** Sankey diagram visualization of the proportion of nuclei positive for either the ventral telencephalic marker *FOXG1*, or the eye territory marker *SIX3* (normalized expression > 1) or none of them for from *CHD1L^+/+^* and *CHD1L^-/-^* hCO. Percentage of nuclei are indicated for each category.

According to our transcriptomic and cistromic data, we reasoned that if CHD1L cooperates with proteins involved in brain development and regionalization, organoids lacking *CHD1L* should exhibit forebrain cell-fate determination defects. To test this possibility and to determine which cell populations are the most affected by the absence of CHD1L, we employed Single Nuclei Gene Expression and ATAC-seq Multiome (referred as snMultiome) analysis to simultaneously profile the transcriptional and chromatin states of control and *CHD1L* mutant (Line 1) 60 DIV organoids. A total of 4,019 nuclei were individually sequenced, including 2,385 nuclei extracted from control organoids, and 1,634 nuclei from *CHD1L^-/-^* mutant organoids (**Figure 7E**). To our surprise, we observed almost no overlap of the nuclei identity clusters from control and *CHD1L^-/-^* mutant organoids showing a dramatic effect of the absence of *CHD1L* on cell identity (**Figure 7E and F**).

The Seurat weighted nearest neighbor method was used to compute a neighbor graph which was visualized with UMAP. In control human organoids, a total of 16 clusters were annotated based on expression of marker genes (**Figure 7G-I and Supplementary Figure S9B**). We found groups with telencephalic identity (FOXG1, PAX6, SOX2) including radial glial cells, cycling radial glial cells, intermediate progenitor cells (IPC), and two groups of neurons including immature and excitatory neurons. We also identified groups with retinal identity (SIX3, RORB, VSX2, OTX2) including Retinal Progenitor Cells, Cycling Retinal Progenitor Cells, Transient Retinal Precursor Cells, Retinal Ganglionic Cells, Amacrine/Horizontal cells and Photoreceptors/Cones. We further found cell clusters from Forebrain Telencephalic/Diencephalic boundary, Mesenchyme, Hindbrain and Choroid Plexus (**Figure 7G-I**).

Strikingly, we observed major differences between control and *CHD1L* mutant organoids. We detected only 11 clusters in mutant organoids mainly due to the absence of the telencephalic clusters (i.e. radial glial cells, cycling radial glial cells, IPC, immature and excitatory neurons). More specifically, we observed that among the 2,385 sequenced nuclei from *CHD1L^+/+^* cerebral organoids, a total of 1,436 nuclei were *FOXG1*-positive (60.2%) whereas 169 nuclei were *SIX3*-positive (7.1%). To the contrary, we found only three *FOXG1*-positive nuclei (0.1%) among the 1,634 nuclei and a total of 809 *SIX3*-positive nuclei (49.5%) for *CHD1L^-/-^* (Line 1) cerebral organoids (**Figure 7J**). These findings exemplified the profound cell fate difference between control and *CHD1L* mutant organoids, the latter expressing retinal-specific genes and resembling to mature hiPSC-derived retinal organoids at 60 DIV(119, 120) (**Figure 7B**).

Based on our snMultiomic data, we thus speculated that *CHD1L* could be a driver regulator for cell-fate decision in the forebrain. To test this possibility, we performed SCENIC+ workflow to catalog the set of enhancer-driven regulons that form gene regulatory networks (GRNs) in both wildtype and *CHD1L* mutant organoids(63) (**Figure 8A and B**). SCENIC+ identified 55 activator and 12 repressor eRegulons (**Figure 8C**). SCENIC+ recovered well-known master regulators of excitatory neurons (NEUROD2, MEF2C, TBR1 and FOXG1), retinal ganglionic cells (RAX, CRX, RXRG, NEUROD1), transient retinal precursor cells (RFX2, MITF) and photoreceptors/cones (ISL1, POU2F2, EBF1/3). The majority of the top five cell-type-specific transcription factors showed co-binding to shared enhancers (**Figure 8D and Supplementary Figure S9C and D**). Individual eRegulon visualization further confirmed the presence of transcription factors favorizing telencephalic fate [FOXG1(+), NEUROD6(+)] and the subsequent positive regulation of their associated genes and regions in *CHD1L*^+/+^ organoids whereas transcription factors associated with retinal fate specification [OTX2(+), VSX2(+), SIX3(+)] were found in *CHD1L* mutant organoids. Of note, the eRegulon SOX2(+) was found in both control and *CHD1L* mutant organoids as expected (**Figure 8E and F and Supplementary Figure S9E**). Taken together, these data revealed that CHD1L acts as a master regulator of cell-fate decision and promotes telencephalon fate during forebrain regionalization.

**Figure 8:**
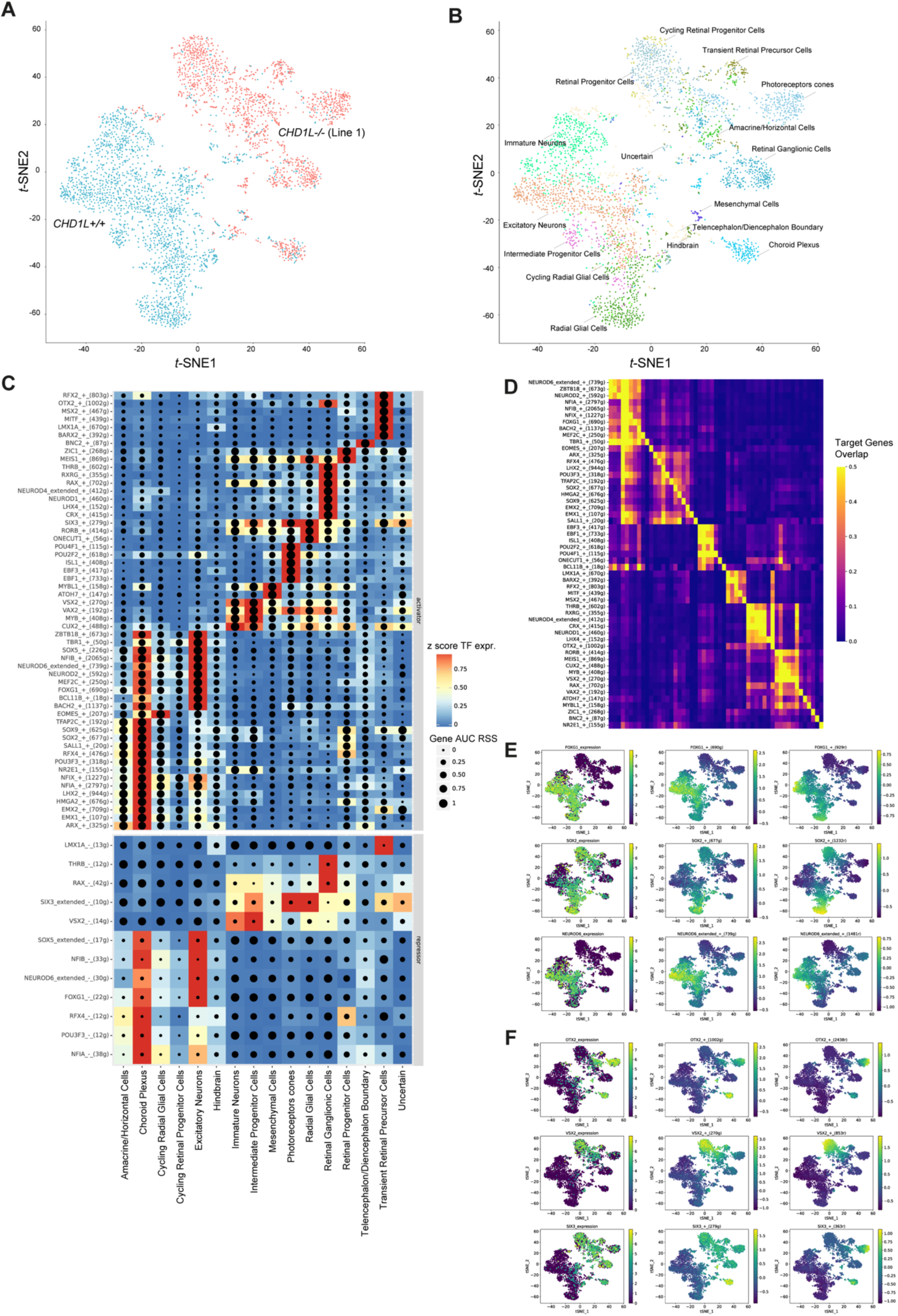
SCENIC+ analysis reveals CHD1L function in forebrain cell-fate determination. **(A)** *t*-SNE dimensionality reduction of 4,019 cells based on gene and target region enrichment scores of eRegulons. Cells are colored according to their genotypes *CHD1L*^+/+^ and *CHD1L*^-/-^ and were analyzed for gene expression and chromatin accessibility in 60 days *in vitro* hCO. **(B)** *t*-SNE dimensionality reduction of organoid cells based on gene and target region enrichment scores of eRegulons. Cells are colored according to their identity. **(C)** Heatmap/dot-plot showing transcription factor (TF) expression of the eRegulon on a color scale and cell-type specificity (RSS) of the eRegulon on a size scale. Cell types are ordered on the basis of their gene expression similarity. **(D)** Overlap of target genes of eRegulons. The overlap is divided by the number of target genes of the eRegulon in each row. **(E)** Representative *t*-SNE dimensionality reduction of multiome datased colored based on TF expression and target gene and region activity of eRegulons for telencephalic markers (FOXG1, SOX2, NEUROD6). **(F)** Representative *t*-SNE dimensionality reduction of multiome datased colored based on TF expression and target gene and region activity of eRegulons for retinal markers (OTX2, VSX2, SIX3). AUC, area under the recovery curve; g, gene; r, region; RSS, eRegulon specificity score; TF, transcription factor.

### ATPase activity of CHD1L is not required to rescue neuroanatomical and body length phenotypes

Human *CHD1L* has two protein-coding isoforms characterized by alternative N-termini: *CHD1L-202* (ENST00000369258.8; 897 amino-acids), encoding a full-length (FL) protein isoform (hereafter referred as CHD1L-FL), and *CHD1L-203* (ENST00000369259.4, 693 amino-acids), lacking the first ATPase lobe (hereafter referred as CHD1L-ΔLobe1), which relevance to the investigation was due to its higher expression in brain-related tissues, as reported in GTEX (https://gtexportal.org/home/gene/CHD1L) and its detected expression in our hiPSC-derived hNPC (**Figure 9A and B**). A functional role for CHD1L- ΔLobe1 isoform is unexpected since a construct of CHD1L lacking its first ATPase lobe has been shown to be devoid of an ATPase activity(32).

**Figure 9:**
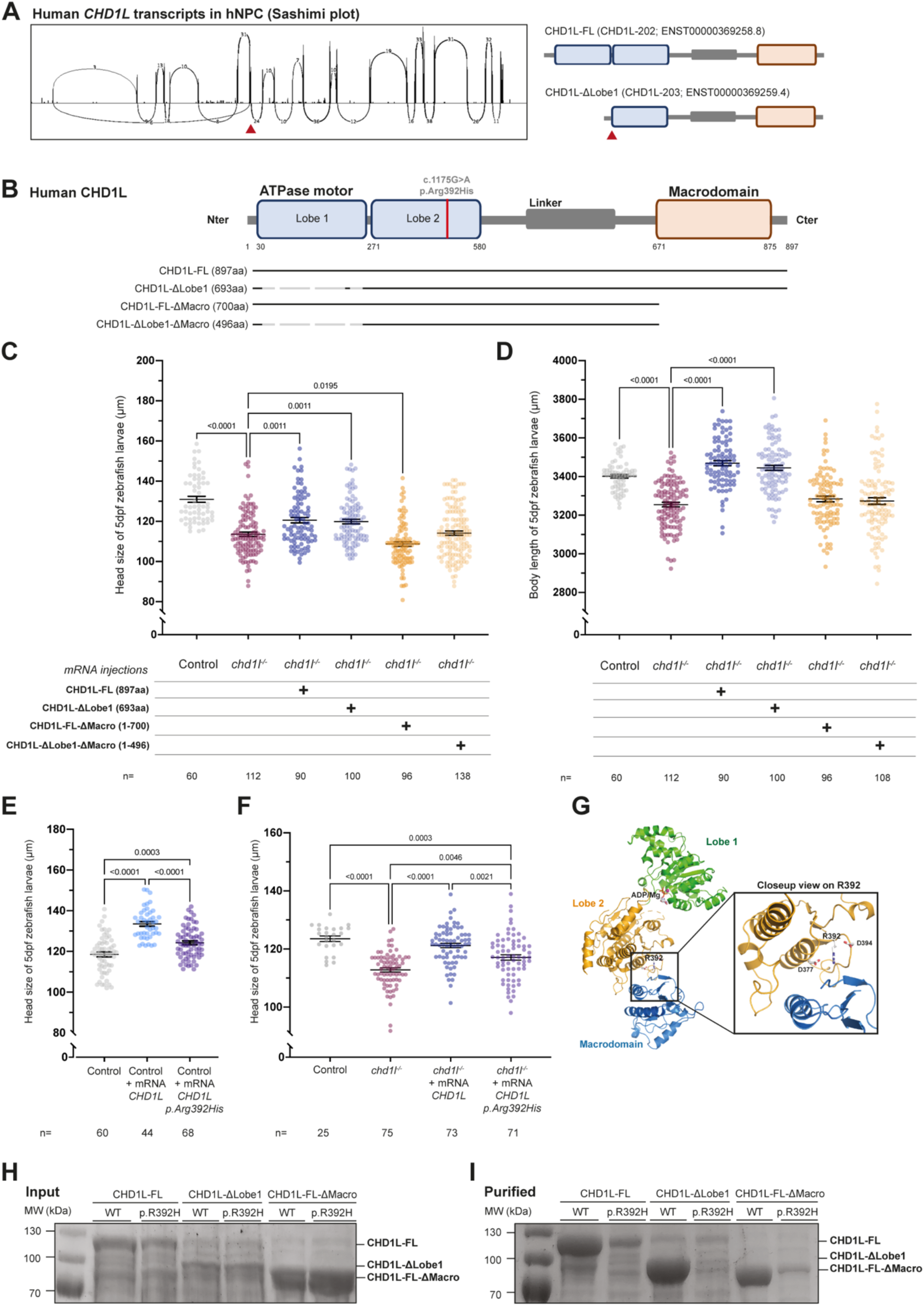
CHD1L function is mediated through its macrodomain and is implicated in neurodevelopmental disorders. **(A)** Sashimi plot of *CHD1L* mRNA transcripts expressed in hNPC showing two main transcripts referring to *CHD1L-202* (ENST00000369258.8) and *CHD1L-203* (ENST00000369259.4). **(B)** Schematic representation of the human protein isoforms of CHD1L including CHD1L-FL (CHD1L-202), CHD1L- ΔLobe1 (CHD1L-203) and the two truncated forms of CHD1L (CHD1L-FL-ΔMacro and CHD1L-ΔLobe1- ΔMacro). The homozygous variant p.Arg392His is shown on the protein. **(C)** Dot plot showing the distance between the eyes (head size) of 5 dpf control larvae and larvae injected with mRNA for each of the two isoforms and the two truncated forms of *CHD1L*. Data shown as mean ± SEM of triplicate batches; Kruskal-Wallis test performed between *chd1l^-/-^* alone versus all the other conditions. **(D)** Dot plot showing the body length of 5 dpf control larvae and larvae injected with mRNA for each of the two isoforms and the two truncated forms of *CHD1L*. Data shown as mean ± SEM of triplicate batches; Ordinary One-Way ANOVA performed between *chd1l^-/-^* alone versus all the other conditions to validate phenotypic rescue. **(E)** Dot plot showing the distance between the eyes (head size) of 5 dpf control larvae or larvae injected with human *CHD1L-FL* mRNA either wild-type or carrying the mutation p.Arg392His. Data shown as mean ± SEM of triplicate batches; Ordinary One-Way ANOVA. **(F)** Dot plot showing the distance between the eyes (head size) of 5 dpf control, non-injected *chd1l* mutant larvae and *chd1l* mutant larvae injected with the human *CHD1L-FL* mRNA either wild-type or carrying the mutation p.Arg392His. Data shown as mean ± SEM of triplicate batches; Ordinary One-Way ANOVA. **(G)** Ribbon representation of the structure of the self-inhibited form of CHD1L (PDB entry 7epu) in full (left panel) and with a closeup on the region harboring Arg392 residue (boxed right panel). Arg392 structures a specific region of CHD1L ATPase lobe 2 (orange) by interacting with main chain carbonyls and aspartate side chains. This region is key in interacting with the CHD1L macrodomain (blue) in the inhibited form. **(H)** Total protein expression levels in *E. coli* of the three constructs used in this study. No significant changes are observed between the wild-type (WT) and p.Arg392His constructs. **(I)** Levels of the soluble CHD1L constructs from (H). The p.Arg392His mutation causes a major decrease of the solubility of CHD1L constructs, irrespective of the construct considered.

We thus sought to determine whether the ATPase activity of CHD1L is necessary during neurogenesis by injecting the *CHD1L-ΔLobe1* mRNA into zebrafish eggs. Similar to the effect of an overexpression of the *CHD1L-FL* mRNA, we observed macrocephaly and increased body length upon injection of the *CHD1L-ΔLobe1* mRNA (**Supplementary Figure S10A-D**). Likewise, overexpression of the *CHD1L-ΔLobe1* isoform was also able to rescue both reduced head size and body length of *chd1l*^-/-^ mutant zebrafish larvae (**Figure 9C and D**). In addition, we explored the consequences of the ablation of the macrodomain in both *CHD1L* isoforms: *CHD1L-FL-ΔMacro* and *CHD1L-ΔLobe1-ΔMacro* (**Figure 9B**). None of these truncated forms was able to rescue the smaller head and body size of the *chd1l^-/-^* mutant zebrafish larvae. (**Figure 9C and D**). Strikingly, these *in vivo* data indicated that the macrodomain but not the ATPase activity is essential to CHD1L’s role during brain development and growth in a vertebrate organism, further supporting the possibility that the remodeling activity of CHD1L is not required during these developmental processes.

### *CHD1L* contribution to neurodevelopmental phenotypes

Finally, we examined whether loss of *CHD1L* in humans might be sufficient to cause some of the commonly observed phenotypes associated with the 1q21.1 distal region deletion. During our analyses, a 260 kb deletion in 1q21.1 encompassing *CHD1L* only that segregated with ASD was discovered in the frame of a large autism cohort study (SSC proband 12719.p1)(121). By interrogating the Decipher database (deciphergenomics.org), we found two heterozygous atypical deletions and one heterozygous atypical duplication affecting *CHD1L* only that segregated with autism and other neuropsychiatric traits. Furthermore, additional three atypical duplications encompassing either *CHD1L* and *FMO5* or *CHD1L* and *BCL9* are associated to complex neurodevelopmental disorders (**Supplementary Figure S11A and Supplementary Table S6**).

In ClinVar database, we found two unrelated individuals carrying a heterozygous mutation in *CHD1L* leading to a stop at the C-terminus of the protein (c.1929del; p.Arg643Serfs*16); one of the two individuals was reported to present with short stature. Strikingly, we have identified the same heterozygous truncating variant in *CHD1L* in a 44-year-old male with ID and ASD, motor delay, speech delay, seizures, facial features and normal growth parameters. Of note, this individual also carries two other variants of unknown significance in *DLGAP2* (c.1696C>T) and *PDPR* (c.1147G>T). Moreover, a deleterious nonsense *de novo* variant (p.Gln600*) has been found in a proband with sporadic ADHD(122). Remarkably, all these deleterious variants led to a truncation of CHD1L’s macrodomain reinforcing the idea that the macrodomain is necessary for CHD1L’s function during neurogenesis.

Although these findings support the possibility that dosage changes of *CHD1L* might contribute to 1q21.1 distal region deletion/duplication neurocognitive and growth phenotypes, we are cautious at interpreting data from these atypical shorter 1q21.1 CNV and heterozygous deleterious variants. In fact, genetic data on general population suggest that *CHD1L* tolerates heterozygous loss-of-functions (pLI=0; The Genome Aggregation Database (gnomAD)) (**Supplementary Table S6**). We therefore propose that in the context of the 1q21.1 distal deletion/duplication syndromes, the phenotypes cannot be attributed solely to *CHD1L* dosage alteration.

Since *CHD1L* is tolerant to heterozygous loss-of-function, we next investigated whether *CHD1L* could act as a susceptibility gene for neurodevelopmental disorders in a homozygous hypomorphic state. To test this possibility, we searched for *CHD1L* homozygous variants in published exome data. A study focusing on the diagnostic yield of exome sequencing to identify disease genes in consanguineous families described the case of a female patient (ID: ER100167) with mild intellectual disability, microcephaly, muscular hypotonia, rigidity, ataxia, intention tremor, hypopigmented macules, EEG abnormalities(123). The patient carries a homozygous variant in the ATPase Lobe 2 of CHD1L (NM_004284.4, c.1175G>A, p.Arg392His in CHD1L-FL protein; this position corresponds to p.Arg188His in the CHD1L-ΔLobe1 protein). The variant was predicted as probably damaging in PolyPhen-2 (score=1.00 in HumDiv and score=0.99 HumVar) and as affecting protein function in SIFT (score of 0.00, median protein conservation=3.01). Furthermore, it was never reported in homozygous form in gnomAD.

We examined whether the p.Arg392His variant could be pathogenic using our zebrafish model. We overexpressed either *CHD1L-FL* wild-type or p.Arg392His mutant mRNA in wild-type eggs. We found that injection of the mutant mRNA leads to macrocephaly and increased body size; however, the amplitude of the phenotypes was significantly lower compared to the wild-type mRNA effect (**Figure 9E and Supplementary Figure S11B**). We next investigated whether the mutant p.Arg392His mRNA could rescue the phenotypes observed in *chd1l*-/- mutants. The mutant mRNA partially rescued the head size and body measurements of the ENU mutants compared to a full rescue of both phenotypes upon injection of the wild-type mRNA (**Figure 9F and Supplementary Figure S11C**).

Arginine residue at position 392 (Arg392) belongs to the ATPase Lobe 2 of CHD1L and contributes to the structural organization of this second lobe. Specifically, Arg392 participates to the structuring of a specific region of this second lobe which interacts with the CHD1L macrodomain when CHD1L is in its auto-inhibitory form (**Figure 9G**)(36). Arg392, however, is not in direct contact with the macrodomain, suggesting that it mostly plays a structuring role for CHD1L. Importantly, although the replacement of Arg392 by a histidine will keep the positive charge of this residue, the shorter histidine side chain will not be able to retain all interactions made by Arg392, which could alter the stability of this region, or of CHD1L.

We further investigated the impact of the p.Arg392His by introducing this mutation into CHD1L-FL, CHD1L-ΔLobe1, CHD1L-FL-Δmacro constructs. We examined at the expression and solubility levels of these mutants compared to the wild-type (WT) proteins upon expression in *E. coli*. Analysis of the total expression levels showed no significant differences between the WT and all mutant proteins (**Figure 9H**). Strikingly, analysis of the soluble fractions revealed that the p.Arg392His mutation caused a major decrease of the quantity of soluble proteins, irrespective of the construct considered (**Figure 9I**). These data showed the important role of Arg392 not only in structuring the ATPase lobe domain but also at the full CHD1L protein level. We also observed that a fraction of p.Arg392His mutant CHD1L protein remained soluble in our experiments which was in accordance with our zebrafish functional data showing partial phenotypic rescue upon injection of the p.Arg392His variant. Overall, these *in vitro* findings strongly indicated that a small amount of p.Arg392His CHD1L protein could still be functional explaining the hypomorphic deleterious effect seen in zebrafish. Our functional investigations underlined the fact that *CHD1L* homozygous mutations found in cohorts with neurodevelopmental disorders should not be discarded during variant filtration and prioritization, as they may represent candidate risk alleles that require further functional validation.

## DISCUSSION

Investigating the biology of rare but relatively penetrant CNV provides a unique opportunity to understand better brain development and cellular mechanisms underlying increased susceptibility to autism. We and others have found some phenotypic drivers for several pathogenic CNV such as 15q13.3 deletions and duplications(124–128), 16p11.2 deletions and duplications (BP4-BP5 and BP2- BP3)(68, 70); and 17q12 deletions and duplications(129) all of which are associated with increased risk for developmental and neuropsychiatric disorders.

Here, we investigated the 1q21.1 CNV-prone locus, the second most common region associated with ASD for which contribution of genes within the distal locus remained to be explored. By combining *in vivo* modeling utilizing zebrafish, mouse, human organoids and omics analyses, our data support a major contributory role for *CHD1L* in the 1q21.1 distal CNV-associated neuroanatomical and growth phenotypes through four lines of evidence: first, the *in vivo* overexpression screen yielded macrocephaly and increased body size in one out of eight genes within the 1q21.1 interval; second, the reciprocal suppression of this gene mirrored the corresponding human 1q21.1 deletion phenotypes; third, omics analyses established neurogenesis impairment that was consistent across species; and fourth, atypical deletions and duplications encompassing *CHD1L* and several deleterious truncating mutations and a pathogenic homozygous *CHD1L* missense mutation further confirmed that *CHD1L* contributes to the neurocognitive phenotypes of 1q21.1 distal deletion/duplication syndrome. In addition to its driver role for brain and growth phenotypes of the 1q21.1 distal CNV, we found that CHD1L acts as a co-transcriptional factor for the pioneer factor SOX2 and CHD1L loss perturbs the neurogenic program in human neuronal progenitor cells. Lastly, modeling the loss of *CHD1L* in self-organizing cerebral organoids revealed that CHD1L is a master regulator of cell-fate decision, specifically favoring telencephalon fate during forebrain regionalization. This work uncovers a novel function for *CHD1L*, extending beyond its well-characterized role in DNA repair and its association with cancer.

Recently, the *NOTCH2NL* paralogs (*NOTCH2NLA* and *NOTCH2NLB*) have been proposed as candidates contributing to the neurocognitive phenotypes of 1q21.1 distal deletion/duplication syndrome. Deletion of these genes leads to premature neuronal maturation whereas ectopic expression leads to a delay in the differentiation of radial glial cells(23, 24). Here, we confirmed that overexpression of *NOTCH2NLB* but not *NOTCH2NLA* led to macrocephaly in zebrafish. In contrast, the simultaneous overexpression of both *NOTCH2NLA/B* did not affect zebrafish head size. This observation prompts us to speculate that *NOTCH2NLB* contributes to the brain size phenotypes associated with 1q21.1 distal CNV, but only when its copy number is increased relative to *NOTCH2NLA*. Reported atypical 1q21.1 CNV support this possibility; Fiddes *et al.* found that *NOTCH2NLB* was completely duplicated in all atypical duplication cases (which were all macrocephalic) and entirely deleted in all atypical deletion cases (which were all microcephalic)(23). Further, we observed that overexpression of either *NOTCH2NLA* or *NOTCH2NLB* had no impact on body size. Taken together, we suggest that the 1q21.1 distal CNV follows a *cis*-epistasis complex CNV model(20) for which multiple primary drivers are sufficient to cause independent or same phenotypes and modifiers are present to modulate the expressivity or penetrance of the phenotypes. Specifically, *CHD1L* and *NOTCH2NLB* dosage changes cause head size phenotypes and autistic traits; *CHD1L* dosage changes cause growth phenotypes; *GJA5* overexpression is responsible for cardiac defects(130); and *GJA8* is responsible for eye defects(18, 19) with *CHD1L* acting as a possible modifier according to our data obtained in human cerebral organoids.

Considering the variable penetrance and phenotypic expressivity, we are under no illusion that other genes outside the 1q21.1 distal region might also contribute to the associated phenotypes. For instance, it has been proposed that 3D proximity of gene loci supports the notion of a co-regulation mechanism of genes with related function(131). One can think about possible co-regulation of autism genes or CNV in -*cis* and -*trans* at the chromatin level. Of note, recent chromatin conformation analysis showed that the 16p11.2 phenotypic modifiers *MVP* and *MAPK3* have long-range chromatin interactions with *PTEN* and *CHD1L*, respectively(132). Analogous to 1q21.1 distal CNV, deletions and duplications of the 16p11.2 BP4-BP5 interval are linked to macro- and microcephaly respectively(68). Regulatory chromatin loops between ASD susceptibility loci/regions should be thus further explored to characterize better the relationship between ASD driver genes, their cellular functions in time and space, and how their modulation affects the penetrance and expressivity of the human phenotypes.

Human growth is a highly complex and multifactorial trait. Rare CNV including the 1q21.1 deletions are a relatively common cause of short stature(14). Our zebrafish data indicate that larval body size can be modulated by *CHD1L* dosage changes only and we found that several genes associated with short stature in humans are downregulated in *CHD1L* mutant hNPC. Remarkably, stable *Chd1l* mutant mice also exhibit decreased body size and decreased body weight(133). Moreover, the height associated SNP rs6658763 has been found in linkage disequilibrium with two *CHD1L* non-synonymous variants (r^2^=0.907) in a large genome wide association study on human adult height(80) which further support the possibility that *CHD1L* influences metabolism and growth. Additional experiments will thus be required to determine how CHD1L controls these biological processes, and which cells are sensitive to *CHD1L* dosage during development.

CHD1L belongs to the SNF2 superfamily(134) and is a poly(ADP-ribose) and ATP-dependent remodeler, with a role in chromatin relaxation. It consists of a two-lobed catalytic Snf2-like ATPase domain, which is connected through a linker region to a C-terminal macrodomain, which mediates PARP1 activity-dependent chromatin-targeting(28–32). Its involvement in the early cellular response to DNA damage was elucidated, providing evidence to the presence of an auto-inhibitory mechanism caused by the interaction of the CHD1L’s macrodomain and the bilobate ATPase module(28–35). DNA-damage-mediated PARP1 induction suppresses this interaction, allowing release from auto-inhibition, activation and therefore chromatin decompaction(36, 134). Our proteomic data confirmed that BER factors and PARP1 are protein partners of CHD1L in hNPC. Published data on adult *Chd1l*^−/−^ mice excludes DNA repair defects as a potential cause for the increased apoptosis observed in our zebrafish experiments; *Chd1l*^−/−^ mice exhibit neither elevated levels of DNA damage across various tissues nor a shortened lifespan, suggesting that DNA repair deficiencies are not responsible for the apoptotic changes we observed in zebrafish larvae(133). The fact that a *Chd1l* mutant murine model is not lethal contradicts a previous study reporting that *Chd1l* was necessary for embryo preimplantation(37). Another study reports that Chd1l has a role during cell reprogramming during the initial step of cell reprogramming in mice, requiring both its macrodomain and DNA helicase activity(135). Strikingly, our study contradicts the necessity of CHD1L’s DNA helicase activity. The presence of a functional CHD1L isoform lacking the first ATPase lobe in humans, along with our zebrafish data, strongly suggests that CHD1L’s remodeling activity is dispensable for brain development and body growth *in vivo*.

To date, *CHD1L* is best known for his role during tumorigenesis(135, 136). More recently, CHD1L has been associated with HIV-1 replication rate(137). In rare diseases, to our knowledge, single nucleotide variants of *CHD1L* have not been linked to neurodevelopmental disorders, possibly due to its high tolerance to heterozygous loss-of-function preventing clinicians to consider *CHD1L* heterozygous variants as relevant in complex neurodevelopmental disorders. Based on our modeling data, *CHD1L* homozygous missense variants should be considered as susceptibility alleles in the future. Heterozygous missense mutations in *CHD1L* have been associated with congenital anomalies of the kidneys and urinary tract also known as CAKUT(138); this is in accordance with our transcriptomic data indicating that CHD1L regulates genes involved in kidney/nephron development. In addition, we recently showed that hypermethylation at 1q21.1 locus influences *CHD1L* dosage and constitutes a susceptibility to multiple sclerosis; the loss of *chd1l* leads to oligodendrogenesis impairment and reduced axonal tract projections in zebrafish(139), which reflects that CHD1L has the potential to control cell fate of various neural lineages.

A common feature seen in cellular models of autism is synaptic defects (e.g. SHANK3 and FMR1)(140–143). Several cellular studies looking at cellular phenotypes of other CNV such as 2p16.3/*NRXN1*, 15q13.3, 16p11.2 and 22q11.21 have also shown synaptic dysfunction(128, 144, 145). Our transcriptomic data revealed that loss of CHD1L leads to decreased expression of genes involved in biological processes such as “neuron differentiation” and “synapse assembly” in hNPC suggesting that *CHD1L* dosage affects synaptic function. In fact, through knockdown experiments in hiPSC-derived neurons, we recently demonstrated that *CHD1L*-deficient neurons exhibited branching abnormalities whereas synaptic density appeared unaffected. Additionally, *CHD1L* knockdown led to reduced calcium signal intensity in neurons, which suggests diminished electrical activity in neurons with lower *CHD1L* expression(139). Taking together, these *in cellulo* data confirm that *CHD1L* expression is necessary to generate electrically competent neurons and point to impairment of calcium dynamic in *CHD1L*- deficient neurons. Remarkably, Chapman and co-workers showed that human iPSC-derived neurons carrying 1q21.1 distal deletion or duplication also exhibit altered synaptogenesis, differential expression of calcium channels, and aberrant neural network activity(21). It is thus reasonable to speculate that cognitive impairment in 1q21.1 distal CNV carriers are due, in part, to altered synaptogenesis and synaptic dysfunction mediated by *CHD1L* dosage imbalance.

Finally, we explored the notion of regionalization during brain development. During our investigation, several lines of evidence pointed to a critical role of CHD1L during brain regionalization. First, Loss of CHD1L impaired the expression of genes controlled by *NKX2-2*, *ERG*, *OTX2*, *BRD4* and *SOX2*, known transcriptional factors regulating cell fate and regionalization of the brain(107, 108, 114, 115, 146, 147). Second, CHD1L binds the SOX and EGR2 motifs in hNPC and our CUT&RUN data are similar to OTX2 and SOX2 ChIP-seq data. Third, proteomic data showed that CHD1L interacts directly or indirectly with pioneer TFs SOX2, OTX2, PAX6 and components of the NURD complex. Notably, the possibility that CHD1L acts during brain regionalization is supported by the observation of structural differences in brain architecture in individuals with 1q21.1 CNV(148). Combining self-organizing cerebral organoids with single-nuclei genomic technologies, we determined that *CHD1L*-deficient cerebral organoids exhibit cell fate anomalies including enrichment of *SIX3*- and *OTX2*-positive nuclei and detection of nuclei clusters such as retinal precursor cells and photoreceptors at the expense of *FOXG1*-positive cells such as excitatory neurons normally expected in cerebral organoids. Strikingly, loss of function of SOX2, OTX2, PAX6, leads to eye abnormalities in humans such as microphthalmia (reduced eye size), anophthalmia (no eye formation at all) and coloboma(149). Similarly, some individuals with 1q21.1 distal deletion exhibit abnormally small eyes, coloboma and lens abnormalities. These eye abnormalities could be thus attributed to *CHD1L* based on on its physical interaction with SOX2, OTX2, PAX6 but also to *GJA8*, for which heterozygous mutations have already been associated with eye phenotypes(18, 150). To our knowledge, interaction between CHD1L and SOX2 in hNPC is novel. Notably, Liu and co-workers have shown that Parp1 stabilizes Sox2 binding to nucleosome which facilitates its pioneer activity at less accessible regions of the genome(151). Therefore, it is reasonable to speculate the existence of a “ménage à trois” involving SOX2, PARP1 and CHD1L at some specific SOX2 motifs, working together to activate the neurogenic program. Further experiments will be needed to test this possibility and to determine whether the ability of CHD1L to bind nucleosomes is necessary in the reported Parp1-Sox2 protein complex in mice.

Overall, we showed that integrating *in vivo* and *in cellulo* modeling of copy number variants across multiple vertebrate species with single-cell technologies is crucial for pinpointing phenotypic driver genes and unraveling the complex interplay between driver and modifier genes. Our functional pipeline is particularly promising for resolving numerous genomic regions implicated in a wide range of human genomic disorder phenotypes. Furthermore, this study highlights CHD1L as a master regulator of cell-fate in the forebrain, along with its identified transcription partners, exemplifying how autism and morphometric traits can result from diverse developmental insults. This underscores the critical importance of characterizing the developmental function of susceptibility autism genes to guide future therapeutic approaches.

## DATA AVAILABILITY & RESSOURCES

RNA-seq, ATAC-seq, CUT&RUN, single-nuclei multiome fastq files have been uploaded to GEO with the accession number GSE266032. All reagent information can be found in Supplementary Table S7.

## SUPPLEMENTARY DATA

Supplementary Figures S1-11 and Supplementary Tables S1-7.

## SUPPLEMENTARY DATA STATEMENT

Data will be available online.

## ACKNOWLEDGEMENTS

We are grateful to Victoria Fischer for her help with zebrafish data acquisition and Mathis Soubeyrand for his technical assistance during organoid production. We thank the IGBMC Zebrafish Facility, in particular Sandrine Geschier for maintenance and care of the zebrafish lines. We thank Nicolas Charlet-Berguerand, Alexandre Reymond and Valérie Schreiber for helpful discussions.

## AUTHORS CONTRIBUTIONS STATEMENT

M.V. Lemée: data curation, formal analysis, investigation, visualization, methodology, and writing— original draft. M.N. Loviglio: data curation, formal analysis, investigation, visualization, methodology, and writing—original draft. T. Ye: formal analysis, data curation, software, visualization, and writing— original draft. P. Tilly: investigation, visualization, formal analysis, writing—original draft. C. Keime: formal analysis, data curation, software, visualization, and writing—original draft. C. Weber: investigation, formal analysis. A. Petrova: investigation, formal analysis. P. Klein: investigation, formal analysis. B. Morlet: investigation, formal analysis, and writing—original draft. O. Wendling: investigation, resources, visualization, formal analysis, and writing—original draft. H. Jacobs: visualization, formal analysis. M. Tharreau: Resources. D. Geneviève: Resources. J. D. Godin: supervision, resources, validation, writing—original draft. C. Romier: supervision, resources, validation, writing—original draft. D. Duteil: formal analysis, visualization, data curation, writing— original draft. C. Golzio: conceptualization, data curation, formal analysis, investigation, supervision, funding acquisition, validation, visualization, methodology, project administration, resources, and writing—original draft.

