## Supplementary Figures for "Disrupted transcriptional networks regulated by *CHD1L* during neurodevelopment underlie the mirrored neuroanatomical and growth phenotypes of the 1q21.1 copy number variant"

**Supplementary Figures S1-11.**

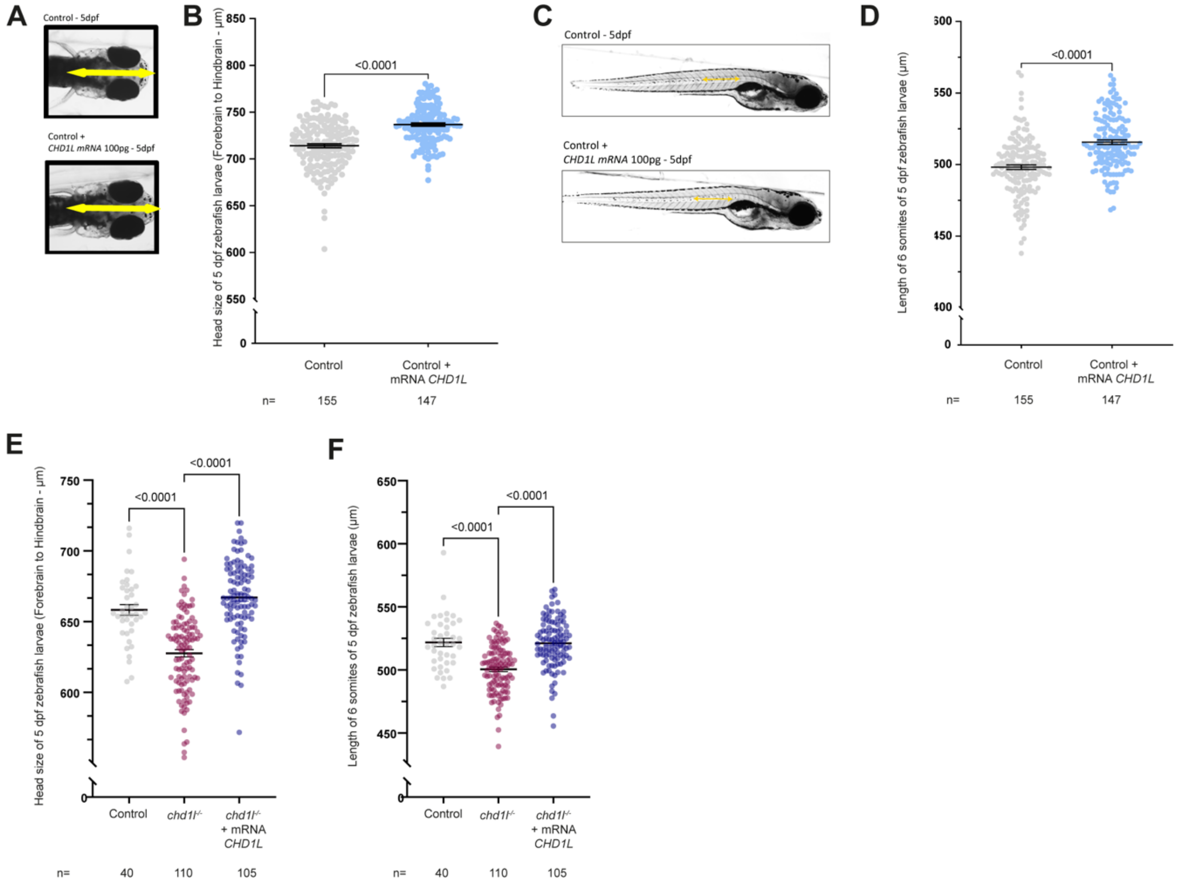


**Supplementary Figure S1: *CHD1L* dosage recapitulates phenotypes associated with 1q21.1 del/dup syndrome**

**(A)** Dorsal views of 5 days post-fertilization (dpf) control larvae and larvae injected with *CHD1L* mRNA, double-ended yellow arrows indicate forebrain/hindbrain measure. **(B)** Dot plot showing the distance between forebrain and hindbrain (head size) of 5 dpf *CHD1L* mRNA-injected larvae compared to control. Data shown as mean ± SEM of triplicate batches; Mann-Whitney test. **(C)** Lateral views of 5 dpf control and *CHD1L* mRNA-injected larvae. Double-ended yellow arrow indicates the 6-somites measurement. **(D)** Dot plot showing the distance between 6 somites of 5 dpf *CHD1L* mRNA-injected larvae compared to control. Data shown as mean ± SEM of triplicate batches; Mann-Whitney test. **(E and F)** Rescue of *chd1*$^{-/-}$ macrocephaly (forebrain/hindbrain distance) and body size phenotypes (6-somites length) by the overexpression of *CHD1L* in *chd1l*$^{-/-}$ larvae compared to control. Data shown as mean ± SEM of triplicate batches; Kruskal-Wallis test. To ease the visualization of the data, the non-significant *p*-values are not showed on the graphs.

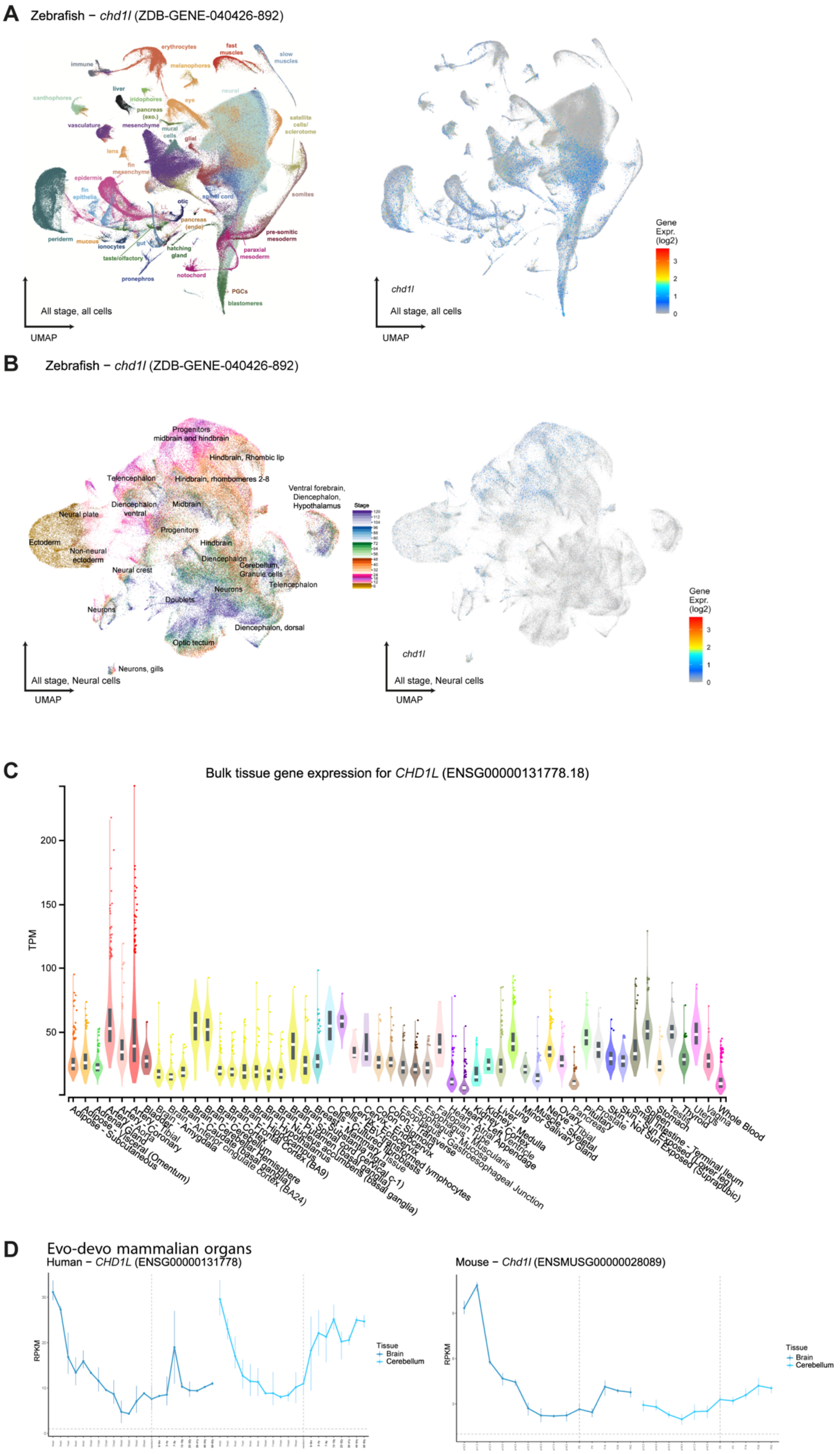
A Zebrafish – chd1l (ZDB-GENE-040426-892)
erythrocytes
fast muscles
slow muscles
immune
melanophores
neural
xanthophores
liver
iridophores
eye
pancreas (exo.)
satellite cells/ sclerotome
vasculature
mesenchyme
mural cells
glial
lens
fin mesenchyme
epidermis
fin epithelia
spinal cord
somites
otic
pancreas (endo)
periderm
mucous
ionocytes
gut
pre-somitic mesoderm
hatching gland
taste/olfactory
paraxial mesoderm
notochord
pronephros
PGCs
blastomeres
All stage, all cells
UMAP
chd1l
Gene Expr. (log2)
B Zebrafish – chd1l (ZDB-GENE-040426-892)
Progenitors midbrain and hindbrain
Hindbrain, Rhombic lip
Telencephalon
Hindbrain, rhombomeres 2-8
Ventral forebrain, Diencephalon, Hypothalamus
Diencephalon, ventral
Midbrain
Neural plate
Progenitors
Ectoderm
Non-neural ectoderm
Hindbrain
Diencephalon
Neural crest
Cerebellum, Granule cells
Telencephalon
Neurons
Doublets
Diencephalon, dorsal
Optic tectum
Neurons, gills
Stage
All stage, Neural cells
C Bulk tissue gene expression for CHD1L (ENSG00000131778.18)
TPM
Adipose - Subcutaneous
Adipose - Visceral (Omentum)
Adrenal Gland
Artery - Aorta
Artery - Coronary
Artery - Tibial
Bladder
Brain - Amygdala
Brain - Anterior cingulate cortex (BA24)
Brain - Caudate (basal ganglia)
Brain - Cerebellar Hemisphere
Brain - Cerebellum
Brain - Cortex
Brain - Frontal Cortex (BA9)
Brain - Hippocampus
Brain - Hypothalamus
Brain - Nucleus accumbens (basal ganglia)
Brain - Putamen (basal ganglia)
Brain - Spinal cord (cervical c-1)
Brain - Substantia nigra
Breast - Mammary Tissue
Cells - Cultured fibroblasts
Cells - EBV-transformed lymphocytes
Cervix - Ectocervix
Cervix - Endocervix
Colon - Sigmoid
Colon - Transverse
Esophagus - Gastroesophageal Junction
Esophagus - Mucosa
Esophagus - Muscularis
Fallopian Tube
Heart - Atrial Appendage
Heart - Left Ventricle
Kidney - Cortex
Kidney - Medulla
Liver
Lung
Minor Salivary Gland
Muscle - Skeletal
Nerve - Tibial
Ovary
Pancreas
Pituitary
Prostate
Skin - Not Sun Exposed (Suprapubic)
Skin - Sun Exposed (Lower leg)
Small Intestine - Terminal Ileum
Spleen
Stomach
Testis
Thyroid
Uterus
Vagina
Whole Blood
D Evo-devo mammalian organs
Human – CHD1L (ENSG00000131778)
RPKM
Tissue
Brain
Cerebellum
Mouse – Chd1l (ENSMUSG00000028089)

**Supplementary Figure S2: *CHD1L* expression across time and species**

**(A)** Single-cell zebrafish transcriptome using UMAP projection for all cell-types and all stages (from 3 hours post-fertilization (hpf) to 120 hpf). Colors show cell identities. Corresponding cells expressing *chd1l* are labelled in blue (Daniocell database). **(B)** Single-cell zebrafish transcriptome using UMAP projection for neural cells from 3 hpf to 120 hpf. Colors show developmental stages. Corresponding cells expressing *chd1l* are labelled in blue (Daniocell database). **(C)** Levels of expression of *CHD1L* in human cell types (GTEx Portal). *CHD1L* is ubiquitously expressed. In brain structures, highest expression of *CHD1L* is observed in Cerebellar Hemisphere (Median Transcript per Million TPM = 55.24), Cerebellum (Median TPM = 52.04), Spinal cord (Median TPM = 40.08). Expression of *CHD1L* is not dependent on gender according to bulk tissue expression (gtexportal.org). **(D)** Expression of *CHD1L* and *Chd1l* in brain and cerebellum during human and mouse development respectively. Highest *CHD1L* expression is observed at 4 weeks post-conception (wpc) (Reads Per Kilobase Million RPKM ≃ 30) and *CHD1L* expression decreases gradually until birth in human cerebrum (RPKM ≃ 5) and cerebellum (RPKM ≃ 8), suggesting an early developmental function of *CHD1L*. Further, *CHD1L* expression increases after birth, mostly in cerebellum (RPKM ≃ 25). This expression pattern before and after birth is conserved in mouse brain structures mostly in the cerebrum.

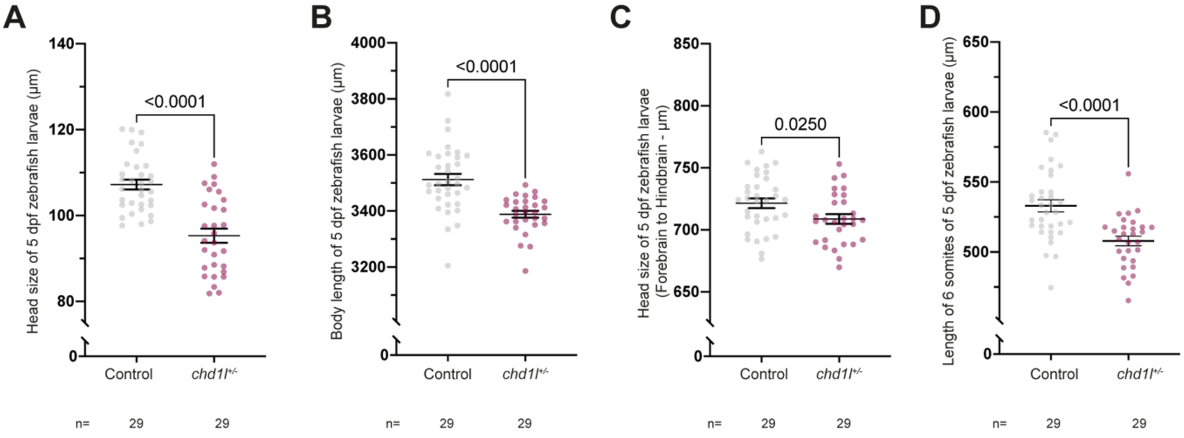


**Supplementary Figure S3: *chd1l+/-* heterozygous zebrafish larvae exhibit same phenotypes as *chd1l-/-* homozygous zebrafish larvae**

**(A)** Dot plot showing the distance between the eyes (head size) of 5 dpf control and *chd1l*$^{+/-}$ larvae. Data shown as mean ± SEM of one of the three replicates; Student's t test. **(B)** Body length of 5 dpf control and *chd1l*$^{+/-}$ larvae. Data shown as mean ± SEM of one of the three replicates; Student's t-test. **(C)** Dot plot showing the distance between the forebrain and hindbrain (head size) of 5 dpf control and *chd1l*$^{+/-}$ larvae. Data shown as mean ± SEM of one of the three replicates; Student's t-test. **(D)** Dot plot showing the distance between 6 somites of 5 dpf control and *chd1l*$^{+/-}$ larvae. Data shown as mean ± SEM of one of the three replicates; Student's t-test.

**A**

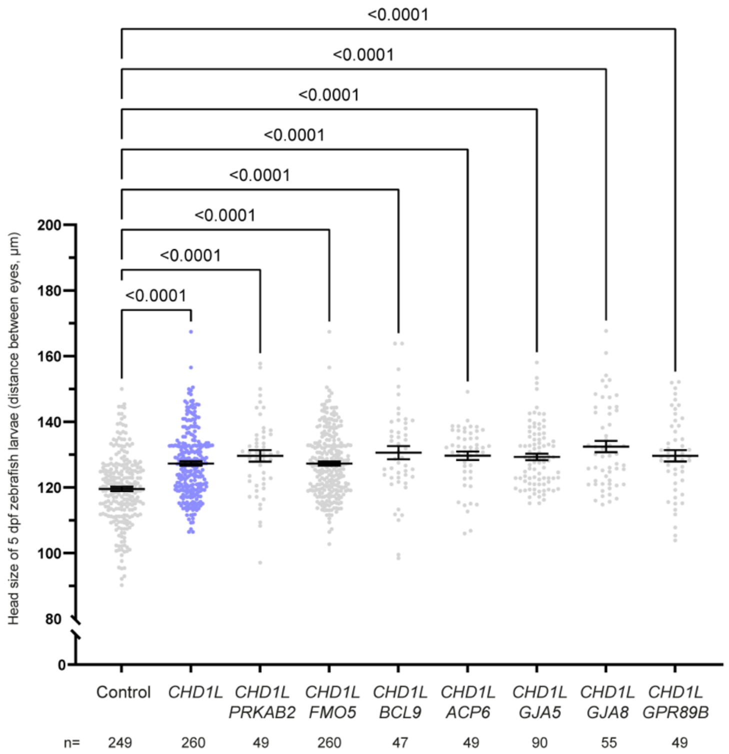


**Supplementary Figure S4: *CHD1L* does not interact genetically with other 1q21.1 distal genes**

**(A)** Dot plot showing the distance between the eyes (head size) of 5 dpf control larvae and larvae injected with *CHD1L* alone or *CHD1L* along with each of the 7 other genes from the 1q21.1 distal region. Data shown as mean ± SEM of at least triplicate batches; Kruskal-Wallis test. To ease the visualization of the data, the non-significant *p*-values are not showed on the graphs.

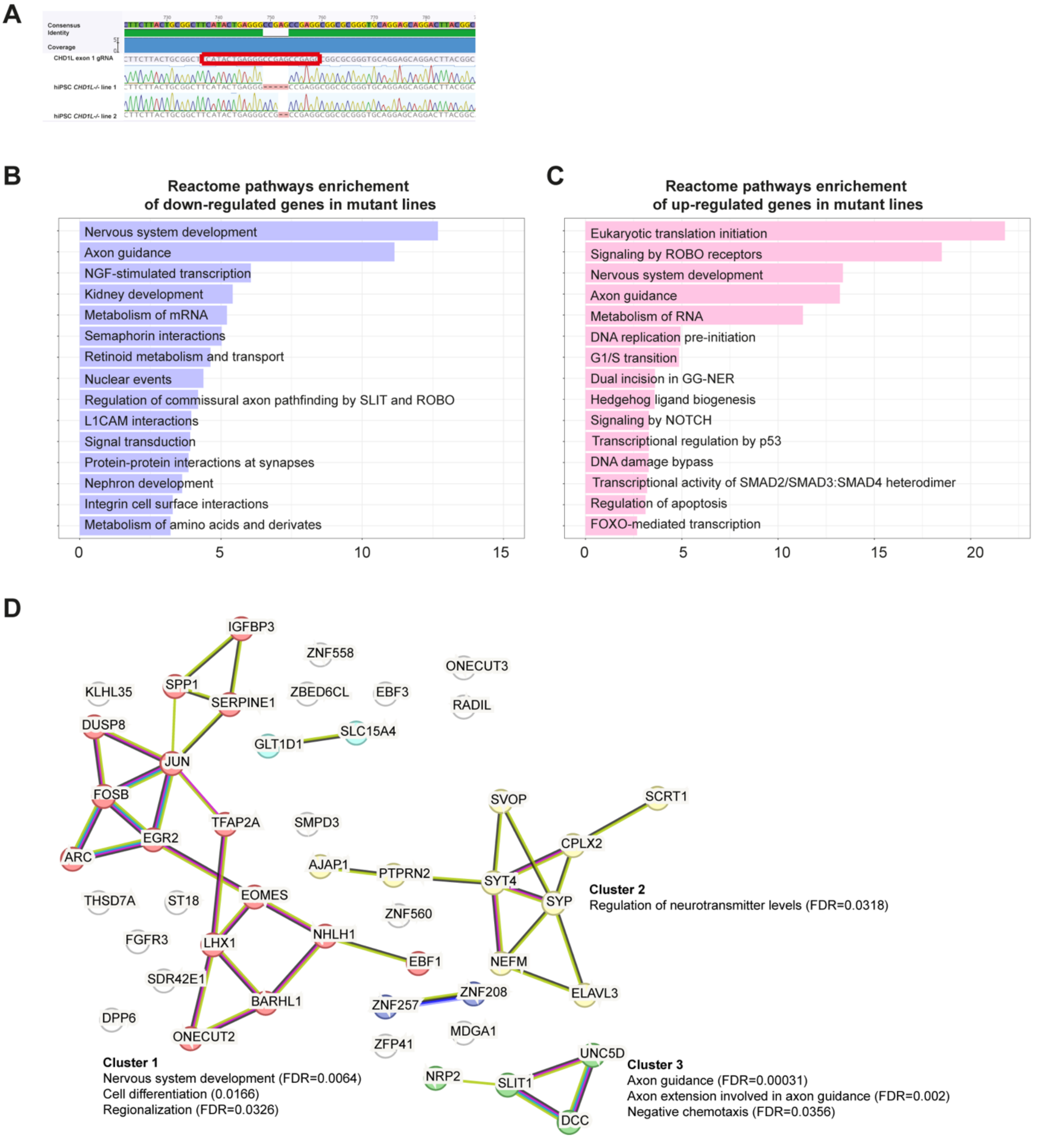


**Supplementary Figure S5: Genome editing at *CHD1L* locus, reactome pathways of all dysregulated genes and STRING network of the 52 top DEG in mutant hNPC**

**(A)** Sequencing of *CHD1L* region in hiPSC *CHD1L*$^{-/-}$ edited by CRISPR-Cas9 shows a 5-base pair (bp) deletion in the first exon of *CHD1L* in mutant Line 1 and a 2 bp deletion in mutant Line 2; both events lead to frameshifts and premature stops. **(B and C)** Reactome Pathways enrichment for the 563 down-regulated and the 294 up-regulated genes in mutant *CHD1L*$^{-/-}$ hNPC respectively. **(D)** STRING analysis of 52 most DEG (|Log2FC|> 1; FDR <0.05) in mutant *CHD1L*$^{-/-}$ hNPC. Proteins are clustered according to k-means and annotated for Biological Process (Gene Ontology). Protein-Protein interactions are showed (PPI enrichment p-value <1e$^{-16}$).

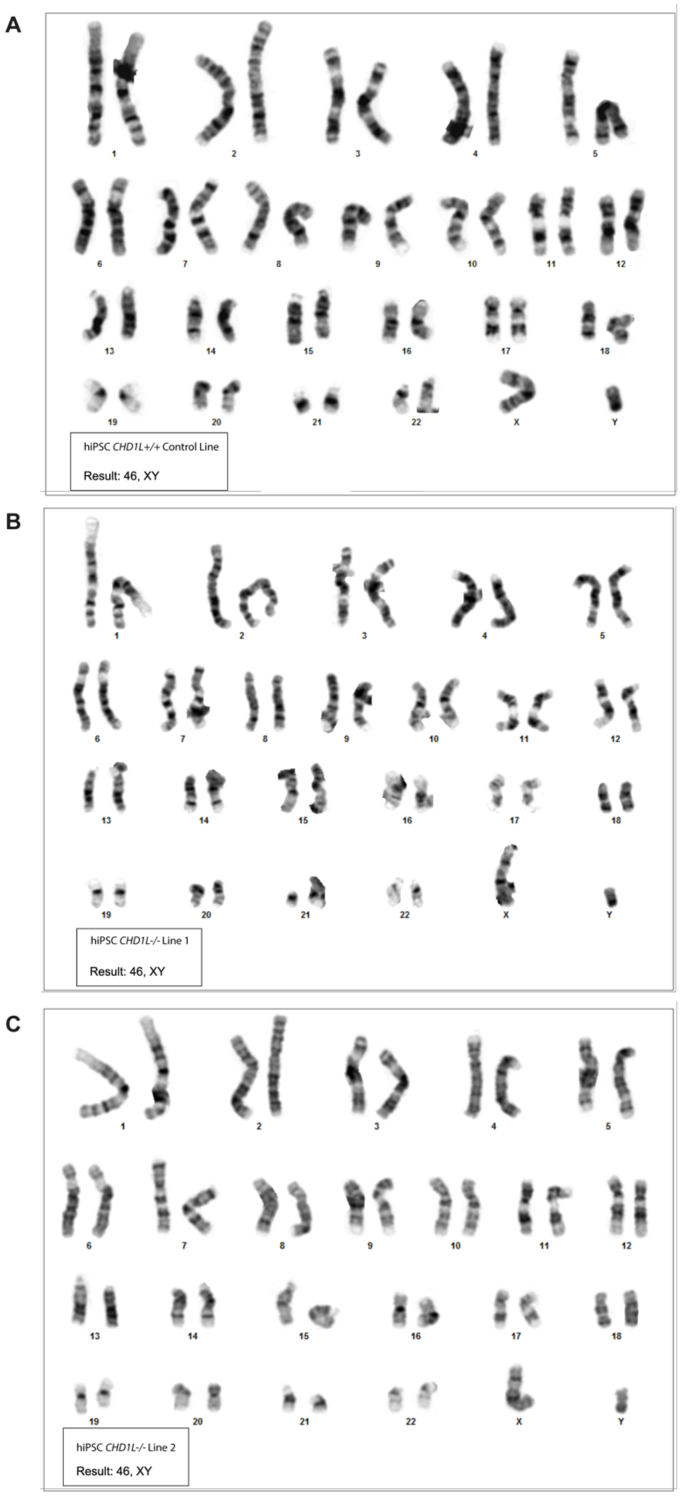


**Supplementary Figure S6: Karyotype of CRISPR-edited and wildtype hiPSC lines**

**(A)** Karyotype of the male hiPSC (GM8330-8) control line. **(B)** Karyotype of the GM8330-8 derived hiPSC *CHD1L*$^{-/-}$ isogenic mutant Line 1. **(C)** Karyotype of the GM8330-8 derived hiPSC *CHD1L*$^{-/-}$ isogenic mutant Line 2.

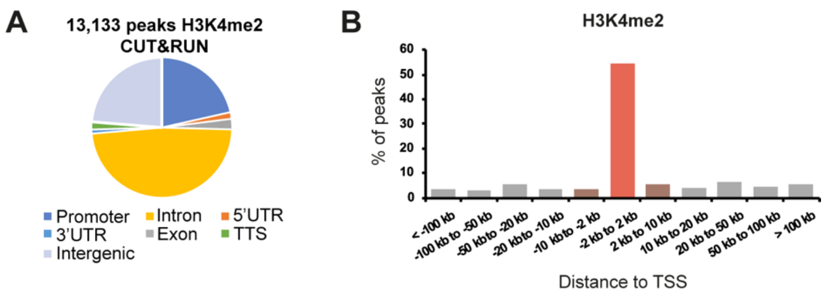


**Supplementary Figure S7: Distribution of H3K4me2 CUT&RUN peaks on chromatin**

**(A)** Pie chart of the distribution of the 13,133 H3K4me2 CUT&RUN peaks on chromatin in hNPC. **(B)** H3K4me2 Peak distribution from transcription start site (TSS). Kilo-base, kb.

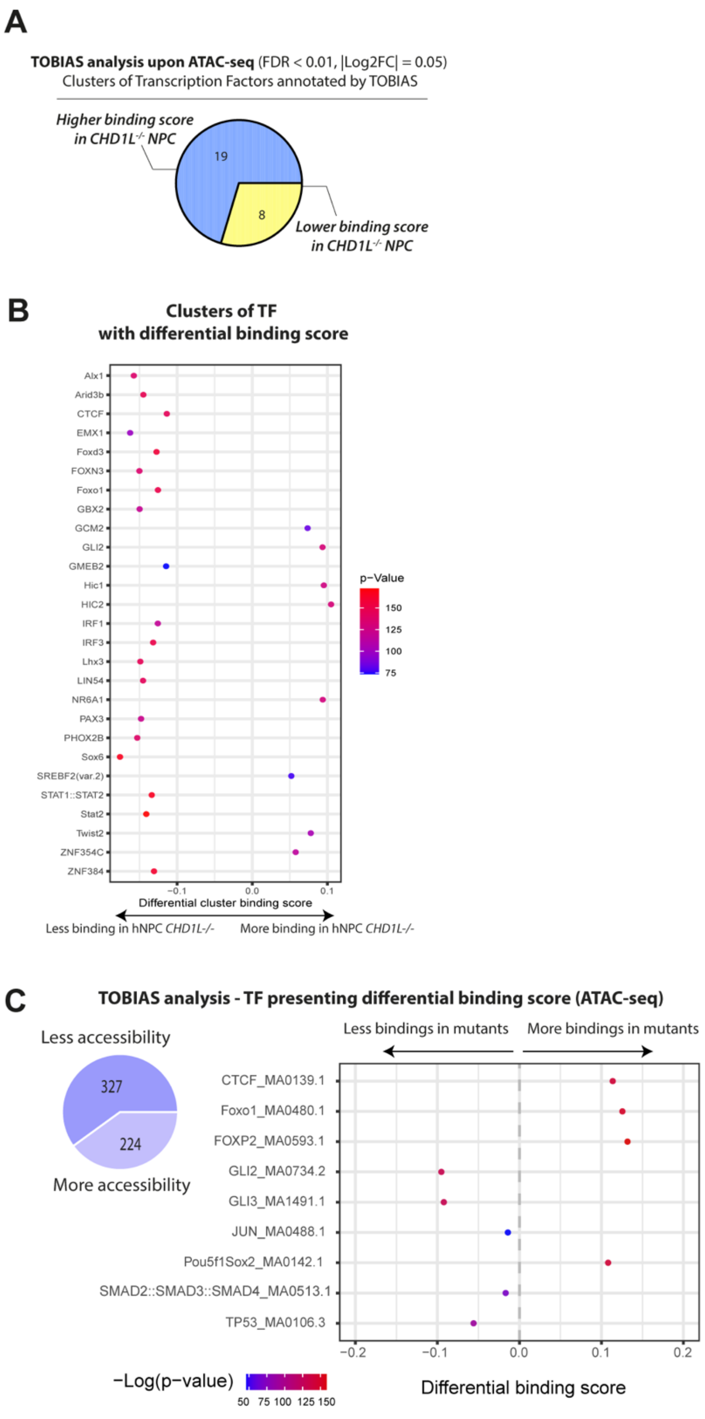


**Supplementary Figure S8: TOBIAS analysis reveals changes in chromatin accessibility of developmental transcription factor clusters**

**(A)** Pie chart of clusters of Transcription Factors presenting differentially binding scores in both *CHD1L*$^{-/-}$ mutant lines compared to control hNPC. Footprint analysis performed using TOBIAS (|Log2FC|> 0.05; FDR<0.01). **(B)** Dot plot of the 27 clusters according to their differential binding scores. **(C)** Example of dysregulated transcription factors bindings predicted by TOBIAS analysis in *CHD1L*$^{-/-}$ hNPC compared to control. TF, transcription factor.

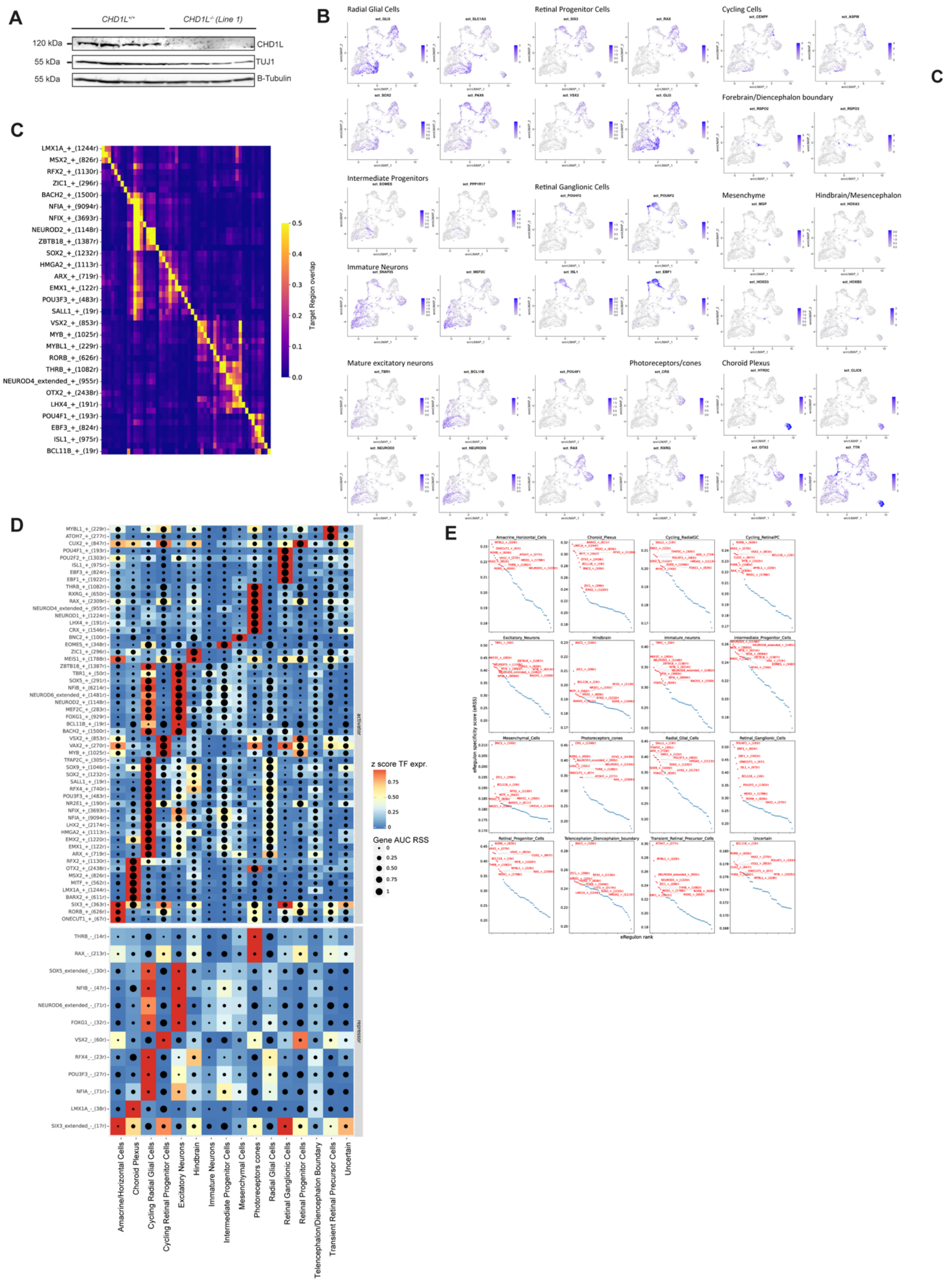


**Supplementary Figure S9: Expression of CHD1L and TUJ1 in organoids, expression of cell markers in snMultiome and eRegulon identified in SCENIC+ analysis**

**(A)** Western Blot of TUJ1 protein level of expression in 52 days *in vitro* $CHD1L^{+/+}$ *and* $CHD1L^{-/-}$ human cerebral organoids (hCO). TUJ1 Expression quantification is shown on Figure 7D. **(B)** wnnUMAP representing expression of cell specific markers in hCO used for cluster annotations. **(C)** Overlap of target regions of eRegulons. The overlap is divided by the number of target regions of the eRegulon in each row. **(D)** Heatmap/dot-plot showing transcription factor (TF) expression of the eRegulon on a color scale and cell-type specificity (RSS) of the eRegulon on a size scale. Cell types are ordered on the basis of their region accessibility similarity. **(E)** Top ten eRegulons assigned to each annotated of hCO cell clusters. AUC, area under the recovery curve; r, region; RSS, eRegulon specificity score; TF, transcription factor.

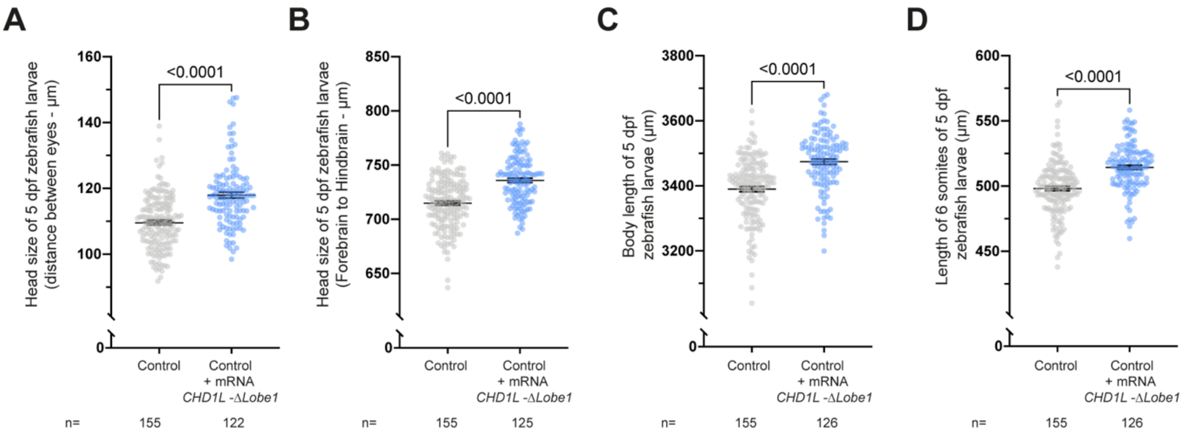


**Supplementary Figure S10: Injection of *CHD1L-ΔLobe1* mRNA induces increased head size and body length similar to the injection of *CHD1L-FL* mRNA**

**(A)** Dot plot showing the distance between the eyes (head size) of 5 dpf control larvae and *CHD1L-ΔLobe1* mRNA-injected larvae. Data shown as mean ± SEM of triplicate batches; Student's *t*-test. **(B)** Dot plot showing the distance between the forebrain and hindbrain (head size) of 5 dpf control larvae and *CHD1L-ΔLobe1* mRNA-injected larvae. Data shown as mean ± SEM of triplicate batches; Wilcoxon test. **(C)** Dot plot showing the body length of 5 dpf control larvae and *CHD1L-ΔLobe1* mRNA-injected larvae. Data shown as mean ± SEM of triplicate batches; Student's *t*-test. **(D)** Dot plot showing the distance between 6 somites of 5 dpf control larvae and *CHD1L-ΔLobe1* mRNA-injected larvae. Data shown as mean ± SEM of triplicate batches; Student's *t*-test.

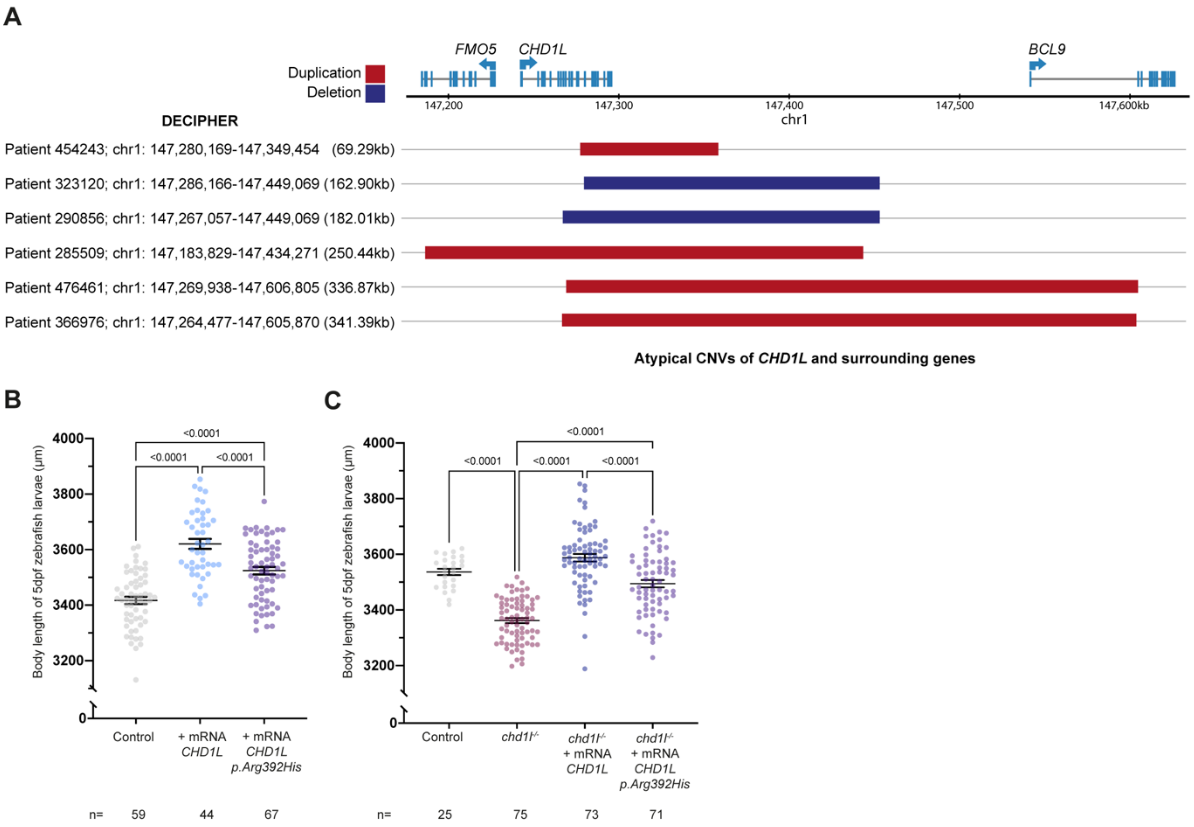


**Supplementary Figure S11: Atypical CNV including *CHD1L* and hypomorphic effect of CHD1L p.Arg392His variant on body length of injected zebrafish larvae**

**(A)** Schematic showing location of atypical copy number variants in 1q21.1 distal region including *CHD1L* alone or along with *FMO5* or *BCL9* (GRCh38) and referenced in deciphergenomics.org. Patient ID, localization and size are indicated. Associated phenotypes are listed in Table 6. **(B)** Dot plot showing body length of 5 dpf control larvae and larvae injected with *CHD1L-FL* mRNA either wildtype or carrying the p.Arg392His variant. Data shown as mean ± SEM of triplicate batches; Ordinary One-Way ANOVA. **(C)** Dot plot showing body length of 5 dpf control larvae, non-injected *chd1l*$^{-/-}$ mutant larvae and larvae injected with *CHD1L-FL* mRNA either wildtype or carrying p.Arg392His variant. Data shown as mean ± SEM of triplicate batches; Kruskal-Wallis test performed between *chd1l*$^{-/-}$ alone versus all the other conditions to validate phenotypic rescue.
